# The biophysical properties of the bacterial nucleoid are dynamic, heterogeneous, and responsive to perturbations of cellular processes

**DOI:** 10.64898/2026.02.18.706702

**Authors:** Xiaofeng Dai, Lauren A. McCarthy, Lindsey E. Way, Emma E. Wiesler, Qin Liao, Anna M. Jensen, Xindan Wang, Julie S. Biteen

## Abstract

Biophysical properties play central roles in cellular function by controlling the diffusion and spatial organization of biomolecules. Because bacteria lack a nuclear membrane, biophysical measurements are often averaged over the entire cell. However, the nucleoid environment is distinct from that of the surrounding cytoplasm, and averaging also ignores local characteristics within the nucleoid. Here, we developed a microrheology framework to quantitatively characterize the bacterial nucleoid and investigate the interplay of its physical properties with cellular processes. We combined single-particle tracking of a genetically encoded protein probe and three-dimensional (3D) Brownian dynamics simulations to separate the nucleoid from the cytoplasm and specifically measure nucleoid accessibility and viscosity. We found that the nucleoid viscosity is 2.5-fold higher than the cytoplasmic viscosity, and that both viscosity and accessibility change systematically across growth phases and the cell cycle. Inhibiting transcription or translation produces opposite changes in nucleoid viscosity in exponential versus stationary phase cells, indicating that the regulation of nucleoid viscosity is sensitive to the underlying biomolecular composition, crowding, and spatial organization. Using Hi-C assays, we further show that changes in nucleoid viscosity may occur without detectable alterations in genome organization, which suggests that nucleoid mechanics provide an independent regulatory mechanism. Spatially, viscosity differences are more pronounced between the nucleoid core and periphery than between genomic locations, and the periphery-core contrast correlates with the coupling of transcription, translation, and membrane insertion. Together, these results indicate that the bacterial nucleoid is a dynamic, heterogeneous viscoelastic environment in which actively regulated biophysical properties may help coordinate multiple cellular processes and provide a physical layer of control that complements canonical biochemical regulation.

**Significance Statement:** The physical properties of intracellular environments strongly influence processes like protein stability and gene regulation. In bacteria, these properties are typically measured across the whole cell. Characterizing the nucleoid environment separately from the chromosome-free region better defines the heterogeneous bacterial microenvironment. Here, we establish an easy-to-implement microrheology framework that combines 2D single-particle tracking of a genetically encoded probe with 3D Brownian dynamics simulations to quantify nucleoid properties, which differ from those of the surrounding cytoplasm and respond to biological processes. Notably, changes in nucleoid viscosity and large-scale genome organization do not necessarily coincide, and nucleoid spatial heterogeneity links to coupling between transcription, translation, and membrane insertion. These findings help resolve the interplay between the physical environment and cellular processes.

## Introduction

The biophysical properties of cellular components regulate a range of essential biological processes, including intracellular transport^1–3^, sub-cellular structures^4–6^, protein synthesis and degradation^7^, and gene regulation and disease^8^. The relatively simple regulatory mechanisms and structural components of bacterial cells suggest a greater reliance on physical principles to optimize and drive biological processes. Accordingly, understanding how physical effects—such as molecular crowding and phase separation—govern intracellular organization and function is crucial^9^. Previous studies have shown that bacterial cells actively expend energy to fluidize their otherwise glass-like cytoplasm, facilitating the mobility and dispersal of large macromolecular complexes^10^. Furthermore, under nutrient limitation, polyphosphate modulates the dynamics of cytoplasmic components in *Pseudomonas aeruginosa*^11^, and the cytoplasmic rheology in *Escherichia coli* is highly correlated with the abundance of amino acid metabolism proteins^12^.

Despite this increasing insight into the relationship between biophysical properties of the cytoplasm and cellular functions, much less is known about the bacterial chromosome, which is condensed into a membraneless organelle called the nucleoid. The nucleoid is a compacted DNA meshwork regulated by nucleoid-associated proteins (NAPs), molecular machines, and enzymes^13^. Previous studies have measured a helical ellipsoid nucleoid shape^14^ and described it as a 50-nm-sized polymer meshwork in a poor solvent^15^. Chromosomal locus motion is consistent with polymer dynamics in a viscoelastic environment^16^, and the nucleoid undergoes rapid structural reorganization in response to perturbations of cellular processes^17^. These findings highlight that the nucleoid physical environment is distinct from that of the surrounding cytoplasm. Given its central role in transcription and translation^18–20^, quantifying the physical properties of the nucleoid is essential for elucidating the interplay between biophysical properties and biological activities.

The physical properties of the subcellular environment can be measured based on the dynamics of intracellular nanoparticles. Techniques including single-particle tracking (SPT)^8,10,11^, fluorescence recovery after photobleaching^3,21,22^, and fluorescence correlation spectroscopy^3,23,24^ have been applied to characterize the cytoplasmic viscosity and its relationship to metabolism and macromolecular organization^25–27^. State-of-the-art microscopy techniques like MPALM^28^ and SMdM^29^ further enable diffusivity mapping. Within the eukaryotic nucleus, such approaches have assessed chromosomal compaction, indicating mesoscale structural changes^30^. In bacteria, probe distributions have been analyzed to study the chromosome structure and its effects on molecular diffusion^15,31^. For instance, Xiang et al. employed probes of varying diameters to quantify size-dependent exclusion from the nucleoid and thereby estimate the average *E. coli* chromosome mesh size^15^. Valverde-Mendez et al. studied intracellular diffusion in 3D to demonstrate the impact of geometrical confinement on ribosome-sized particles and detected distinct diffusive behaviors inside vs. outside the nucleoid, which are largely obscured in traditional two-dimensional (2D) imaging^31^. The nucleoid biophysical properties are important for bacterial physiology. Specifically, viscosity tunes molecular diffusion and thereby diffusion-limited reaction kinetics. In eukaryotes, relatively open euchromatin is enriched in active genes, whereas compact heterochromatin is generally associated with transcriptional repression^32^. Similarly, in bacteria, the nucleoid accessibility determines how readily DNA loci can be reached by regulatory proteins and enzymes. Still, quantitative measurements of these properties have yet to be realized in living bacterial cells, owing to the small size, confined geometry, and membraneless nature of the nucleoid, as well as the absence of accessible, high-throughput methodologies.

To address this challenge, we developed a passive microrheology framework based on tracking a genetically encoded 25-nm fluorescent nanocage probe^33^. This protein assembly is comparable in size to mesoscale biomacromolecules such as the RNA polymerase holoenzyme (10 × 10 × 16 nm)^34^, the 70S ribosome (∼20 nm in diameter)^35^, the replicative DNA polymerases (the PolIII*α*-clamp-exonuclease-*τ*_c_ complex is ∼22.5 nm in its longest dimension)^36^, DNA gyrase (∼22 nm, GyrA₂B₂ heterotetrameric holoenzyme)^37^, and the MukBEF complex (∼24 nm long in its folded state)^38^. By analyzing 2D trajectories acquired using a conventional epifluorescence microscopy setup that is widely available in biology and biophysics laboratories, and then analyzing the data in combination with 3D Brownian dynamics (BD) simulations, we reconstructed the 3D probe diffusion statistically, and we isolated the motion of the probe within the nucleoid from the rest of the cell. This approach enables the direct quantification of the apparent nucleoid accessibility and effective viscosity at the 25-nm scale in living *E. coli* cells.

Our measurements show that the nucleoid is significantly more viscous and sensitive to perturbations than the surrounding cytoplasm. The effective nucleoid viscosity and apparent accessibility change with the growth phases and through the cell cycle. Arresting transcription and translation have opposite effects on the effective nucleoid viscosity in exponential phase vs. stationary phase, which suggests that nucleoid mechanics are mediated by the biomolecular environment. Integration with chromosome conformation capture (Hi-C) data further demonstrates that viscosity changes can occur without detectable alterations of the large-scale genome organization, which indicates that the mechanical and structural states of the chromosome can vary independently. Extending our analysis to the sub-nucleoid level, we uncovered spatial heterogeneity in the chromosomal viscosity: the effective viscosity increases from the nucleoid core to its periphery, and this gradient is modulated by transcription, translation, and inner membrane fluidity. These findings demonstrate that the bacterial chromosome has dynamic and heterogeneous physical properties that can adaptively respond to stress and environmental changes.

This work presents a broadly accessible framework for quantitatively mapping the biophysical properties of the bacterial nucleoid in living cells. The method is compatible with commercial fluorescence microscopes and adaptable to other bacterial species with defined morphologies. By applying our approach to a well-characterized model organism, we determined how cellular processes influence the physical properties of the nucleoid. The bacterial nucleoid is a mesoscale, phase-separated structure^39–42^; characterizing this organelle offers a relevant model for understanding the material properties of chromatin biomolecular condensates more broadly. Furthermore, these measurements provide quantitative parameters for artificial cell design^43,44^ and *in silico* cell reconstruction and modeling^45^.

## Results

### Experimental mapping of probe diffusion combined with Monte Carlo simulations quantifies the apparent nucleoid accessibility

To perform passive microrheology in living cells, we expressed 25-nm sfGFP-labeled protein nanocages^33^ in exponential-phase *E. coli* cells grown at 30 °C in HDA media supplemented with 0.2% glycerol and tracked the particles within the cells at 25 frames per second (Fig. 1A). We use this condition as an example to illustrate our framework for the measurement of apparent accessibility and effective viscosity at the 25-nm-probe scale. We applied this integrated experimental, simulation, and analysis pipeline to all subsequent experiments. Notably, for every condition, we performed custom simulations constrained by the specific cell geometry and optimized parameters through direct comparison with the experimental data for each condition.

**Figure 1.**
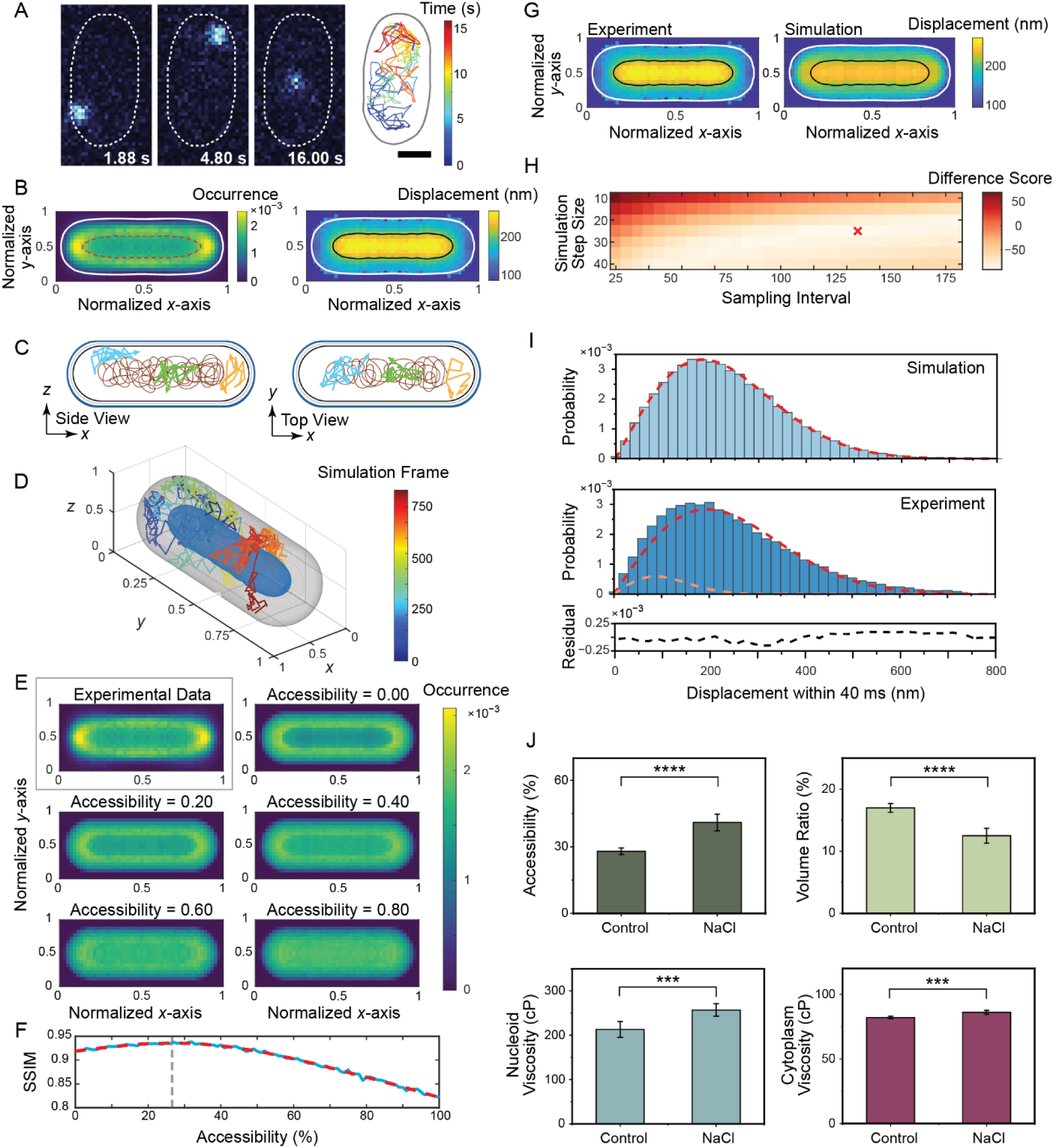
Measurement of the physical properties of the bacterial nucleoid using particle tracking microrheology combined with 3D Brownian dynamics simulations. **A,** Widefield fluorescence microscopy images of an sfGFP-labeled nanocage within a representative *E. coli* cell (left) and a representative nanocage trajectory within the cell (right). Scale bar: 500 nm. The cell outline is derived from segmentation of the phase-contrast image. **B,** Localization heatmap (left) and spatial displacement map (right) from *N* = 2096 cells. Solid white lines: average cell boundary derived from analysis of the heatmap gradient (Methods). Red dashed line: position of the steepest heatmap gradient within the cell, delineating the region inside which nanocages are excluded by the DNA meshwork. Solid black line: smallest displacement map contour line that encompasses the exclusion region identified from the heatmap. **C,** Schematic illustrating the three-dimensional diffusion of the probe within the *E. coli* cell projected onto the *x*-*z* (side view) and *x*-*y* (top view) planes. Probe diffusion is classified into three populations based on compartmental confinement: (1) within the cell poles (orange), (2) within the gap between the nucleoid and inner membrane (blue), and (3) within the nucleoid (green). Brown: illustration of the nucleoid. **D,** 3D Brownian dynamics simulation setup using the reconstructed cell geometry from the heatmap and displacement map in B. Outer and inner confinement boundaries are derived from the white solid and black solid contour lines in B, respectively. A random walk model with rigid collision boundaries was used to simulate probe diffusion within the cell geometry. **E,** Localization heatmaps from experimental data (gray box) and from simulations performed with varying nucleoid accessibility settings. Accessibility is the ratio of the probe density within the nucleoid to the probe density in the cytoplasm in 3D. For each accessibility setting, probes were randomly placed in the reconstructed geometry to generate simulated heatmaps (Methods). **F.** Apparent nucleoid accessibility determined by quantitative comparison of experimental and simulated heatmaps by a structural similarity index measure (SSIM) as a function of simulated accessibility. The SSIM curve (solid blue line) was fit to a polynomial (red dashed line) to find the peak (grey dashed line). **G,** Spatial displacement maps from experiment and simulation optimized by similarity analysis in H. White and black solid lines: cell outline and nucleoid boundary, respectively. **H,** Hyperparameter tuning of 3D Brownian Dynamics simulation by optimization of the difference score between experimental and simulated displacement maps. A composite difference score was calculated using a weighted sum of the *l*^2^-distance to minimize step-size differences and the Pearson correlation coefficient to preserve spatial patterns (Methods). The cross indicates the minimal difference score (and thus the optimal hyperparameter selection). **I,** Distribution of probe step sizes within the nucleoid 2D projection. The cytoplasm simulation step sizes fit a single Rayleigh distribution (top). Experimental step sizes are fit to a two-component Rayleigh distribution (middle), consistent with one component matching cytoplasmic diffusion as in the simulation (red dashed line) and the second capturing diffusion within the nucleoid (orange dashed line). Component weights are calculated from the apparent nucleoid accessibility values in F combined with the 3D nucleoid geometry in D. Bottom: residuals of the experimental data fit. **J,** Comparison of cellular physical properties—apparent nucleoid accessibility, nucleoid volume ratio, effective nucleoid viscosity, and effective cytoplasmic viscosity—in untreated *E. coli* cells and cells under osmotic shock induced by 200 mM NaCl. Error bars: 95% confidence interval (CI) from 50 iterations of bootstrap sampling. Two-sample t-tests with Welch correction between the 2 conditions were performed for each measurement (***: *p* ≤ 0.001; ****: *p* ≤ 0.0001).

We identified each cell boundary from the phase-contrast image and built symmetrized 2D histograms of the normalized, super-localized positions of nanocages in *N* > 2000 cells to determine the average localization probability distribution (Fig. 1B, left). We refer to these 2D histograms as heatmaps throughout the manuscript. We also constructed a symmetrized 2D displacement map to determine the average frame-to-frame nanocage displacement as a function of position within the cells (Fig. 1B, right). We determined the cytoplasm boundary (white lines in Fig. 1B) from the inflection point of the gradient of the nanocage localization heatmap. We determined the region where nanocages are partially excluded by the nucleoid (red line in Fig. 1B, left) from the maximum gradient of the localization heatmap (see SI Appendix)^31^. This exclusion differs from the findings of Xiang et al.^15^, likely due to differences in the strains and growth media used, but the average nucleoid periphery identified by this approach is similar to the nucleoid periphery determined by McCarthy et al. based on HUα-PAmCherry super-resolution imaging^46^.

The largest step sizes in the displacement map occur at the cell center region and colocalize well with the area of nanocage exclusion in the heatmap: the solid black contour line in Fig. 1B, right, which fully encompasses the area of partial nanocage exclusion, is at 78% of the maximum displacement. This finding agrees with previous studies that showed protein diffusion is geometrically confined and location-dependent^31,47^, and our simulations of the diffusion of nanocages excluded from a central nucleoid in a rod-shaped cell verify that the confinement of a probe between a nucleoid and an inner membrane can result in this displacement map pattern (Fig. S1A-C). Specifically, the higher probe density in cytoplasmic gaps dominates the 2D projection. Confinement within these gaps produces large apparent displacements at the 40-ms scale. Thus, the average step size is greatest in the cell-center region (Fig. S1).

Because the probe experiences distinct geometric environments within the cell, we separated its diffusion inside the *E. coli* cell into three populations (orange, blue, and green tracks in Fig. 1C, respectively): (1) at the cell poles, which are typically nucleoid-poor; (2) in the gap between the nucleoid and the inner membrane, which constrains cytoplasmic diffusion; and (3) within the nucleoid, where probe motion depends on the nucleoid material properties. In the 2D imaging setup, the projection of 3D trajectories onto the *xy* plane overlaps populations (2) and (3) in the same cell-center region, even though the environments are distinct.

We describe probe motion as a random walk in two distinct environments, the cytoplasm and nucleoid, assuming uniformity within each compartment. Using static and dynamic simulations within reconstructed 3D cellular geometries, our framework estimates the cytoplasmic contribution to the projected cell-center signal and statistically separates it from the nucleoid contribution. The localization heatmap reports the spatial partitioning of the probe between the cytoplasm and nucleoid, from which we infer apparent nucleoid accessibility. This quantity is primarily interpreted as steric exclusion of nanocages by the nucleoid meshwork. The displacement map and displacement distributions reflect the probe motion over the 40-ms frame interval, from which we estimate effective viscosity while accounting for geometric confinement and the 2D projection. These quantities characterize the cellular environment at the spatial and temporal scales sampled by the probe.

The nucleoid accessibility is an important property because it regulates the recruitment of molecules to the nucleoid^15,19^. Here, we sought to measure the accessibility of the nucleoid based on the density of the probe within the nucleoid (population (3)). Therefore, we performed Monte Carlo simulations within a reconstructed cell geometry (Fig. 1D).

We modeled the average cell boundary and the nucleoid boundary in 3D by rotating the corresponding outlines from the displacement maps (white line and black solid lines in Fig. 1B, right) around the long axis, based on the rotational symmetry of the *E. coli* cell and nucleoid (Fig. 1D). We define the apparent nucleoid accessibility as the ratio of the probe density (localization per volume) within the nucleoid to the probe density within the cytoplasm, and we performed Monte Carlo simulations for each accessibility value by randomly positioning probes throughout the cytoplasmic and nucleoid regions with probabilities determined by the accessibility (see SI Appendix). We generated 2D localization heatmaps for each accessibility value from these 3D static simulations (Fig. 1E), and we calculated the similarity between the experimental heatmap and these simulated heatmaps with the structural similarity index measure (SSIM)^48^.

Our simulated heatmaps diverge from the experimental results near the cell boundaries and at the cell poles in the localization heatmap due to averaging over various cell morphologies in the experiments and heterogeneities such as protein aggregates at the cell poles^47,49,50^ that are unmodeled in the simulations (Fig. S1D). However, differences at the endcaps and the cell boundary remain relatively constant with fluctuations, and changes in similarity are driven primarily by the center region (Fig. S1E-G). Therefore, maximizing the similarity between simulated and experimental heatmaps reflects the accessibility of the nucleoid.

We measured the apparent nucleoid accessibility at the 25-nm scale in these exponential-phase cells to be 27% from the maximum of the fit of the SSIM vs. accessibility curve, ensuring a more robust measurement unaffected by noise (Fig. 1F). Overall, this static simulation within the reconstructed 3D cell geometry enables quantification of the apparent nucleoid accessibility while accounting for geometric effects that are inaccessible to conventional 2D measurements^15,46,51^.

### The measurement of effective nucleoid viscosity demonstrates differences in the properties of the nucleoid and the cytoplasm

To decompose tracks from nanocages within the nucleoid from those in the gap between the nucleoid and the inner membrane (Fig. 1C), we simulated nanocage diffusion within the cytoplasm (nucleoid accessibility set to 0 in the reconstructed cell geometry) using a 3D BD simulation based on a random walk model^52^. We simulated probe diffusion by simulating micro-steps that were substantially smaller than the experimentally observed frame-to-frame displacements. At each micro-step, the step length was drawn from a Rayleigh distribution with a prescribed mean, and the direction was chosen randomly in 3D. Boundary crossings were checked at every micro-step, and any step crossing the cell membrane or nucleoid boundary was reflected to keep the simulated probe within the cytoplasm. To obtain displacements comparable to the experimental measurements, we retained probe positions every *N* micro-steps, where the sampling interval represents the 40-ms experimental frame interval. We optimized this cytoplasmic diffusion simulation for agreement with experiments by tuning two key parameters: the simulation step size and the sampling interval. To address the mismatch caused by comparing simulated cytoplasmic probe diffusion to experimental diffusion in the nucleoid and the cytoplasm, we identified the optimal parameters based on a difference score comprised of the *l*^2^-distance in the cytoplasmic region (between the white and black lines in Fig. 1G) and the Pearson correlation coefficient (PCC) over the whole cell area (Fig. 1H and Table S7; see SI Appendix).

Based on the optimal dynamic simulation parameters, we generated a map of simulated displacements (Fig. 1G, right). In this map of cytoplasmic diffusion, the step sizes within the nucleoid region (black line in Fig. 1G) of the 2D projected map refer only to probe motion in the gap between the nucleoid and the inner membrane. We described this motion by fitting the probability density function (pdf) of the simulated step size distribution (Fig. 1I, top) to a Rayleigh distribution, *f*(*d*):

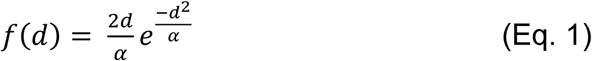

Here, *d* is the displacement per frame time, and *α* is the Rayleigh distribution scale parameter.

On the other hand, the experimental distribution of displacements in the same region is composed of both nanocages in the gap and nanocages within the nucleoid (Fig. 1C). We therefore fit the pdf of experimental step sizes to a two-component Rayleigh distribution (Table S6 and Fig. S2), which is the weighted sum of the Rayleigh distribution from the simulation (Fig. 1I, top) and a second Rayleigh distribution that captures the probe dynamics within the nucleoid (Fig. 1I, middle; see SI Appendix); the weights of these two components are derived from the 27% nucleoid accessibility values calculated in Fig. 1F.

For these exponential phase cells, we found *α* = 1.53 × 10^4^ nm^2^ within the nucleoid from fitting, and we calculated a diffusion coefficient of *D* = 0.096 µm^2^/s for probes diffusing in the nucleoid based on *α* and the experimental imaging frame time, Δ*t* = 40 ms, according to:

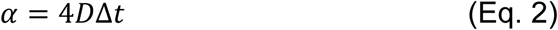

Finally, we calculated the effective viscosity, *η* = 186 cP, within the exponential-phase *E. coli* nucleoid based on the Bolzman constant, *k_B_*, the temperature *T* = 303 K, the radius of the probe, *r* = 12.5 nm, and *D*:

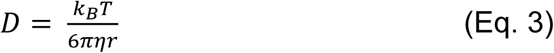

Separately, we characterized the diffusion of probes within the cytoplasm by fitting the experimental distribution of step sizes in the region between the cell inner boundary and nucleoid boundary (white and black lines in Fig. 1G, respectively) to a Rayleigh distribution, and we calculated the cytoplasmic diffusion coefficient and viscosity using Eqs. (2) and (3).

The nucleoid viscosity is significantly higher than the cytoplasm viscosity (213±18 cP vs. 82±1 cP, mean±95% confidence interval from bootstrap sampling, *p* < 0.0001), which demonstrates phase separation between the bacterial nucleoid and the cytoplasm despite the absence of a nuclear membrane^39–42^. This result, which is consistent with the *E. coli* cytoplasm viscosity of ∼100 cP measured by 3D imaging by Valverde-Mendez et al.^31^, highlights the importance of measuring nucleoid viscosity separately from cytoplasmic viscosity rather than characterizing the average bacterial cell viscosity, especially for processes involving DNA^3,16,53,54^.

### Osmotic shock affects the *E. coli* nucleoid and cytoplasm differently

We applied our method to study the effect of osmotic shock with 200 mM NaCl on exponential-phase cells. We measured increases in the effective viscosities of both the cytoplasm and the nucleoid that match the response predicted by polymer physics and rheology (Fig. 1J)^55–59^. The nucleoid viscosity changes more following this perturbation than the cytoplasmic viscosity, which further motivates this approach of measuring the rheological properties of these two subcellular regions separately. The nucleoid also compacts upon osmotic shock (decreased volume ratio), whereas the apparent nucleoid accessibility increases, which indicates that the compacted chromosome meshwork is flexible. Control experiments using SYTOX orange DNA staining further support that the nucleoid compacts significantly upon osmotic shock (Table S8). This phenomenon is consistent with *in vitro* studies of the abundant NAP HU: at high-salt concentrations, HU compacts DNA without stiffening it and dissociates from DNA more rapidly^60^. This compact yet flexible configuration ensures that, though the nucleoid volume is small, the DNA meshwork remains capable of dynamic remodeling *in vivo*. The osmotic shock results are attributed to the effects of physiological osmoadaptation during the 60 min of imaging, rather than an acute osmotic shock response.

### The physical properties of the nucleoid vary with the growth phase and the cell cycle

We measured the *E. coli* effective nucleoid viscosity and apparent accessibility at defined time points along the growth curve (Fig. 2A): at early exponential phase (5 h), at mid-exponential phase with maximal growth rate (10 h), during the transition from exponential phase into stationary phase (15 h), and at stationary phase (24 h). The nucleoid accessibility increases throughout the exponential phase from 5 to 15 h (Figs. 2B, S3A). In stationary phase, however, the nucleoid accessibility decreases (Figs. 2B, S3A), consistent with a shift from biomass accumulation to long-term survival during nutrient limitation^61^. Reduced accessibility is thought to help protect the genome from oxidative and other stresses, while still permitting essential transcriptional and translational activity within the chromosome, through NAPs that reorganize chromosome structure^13,46,51,62^.

**Figure 2.**
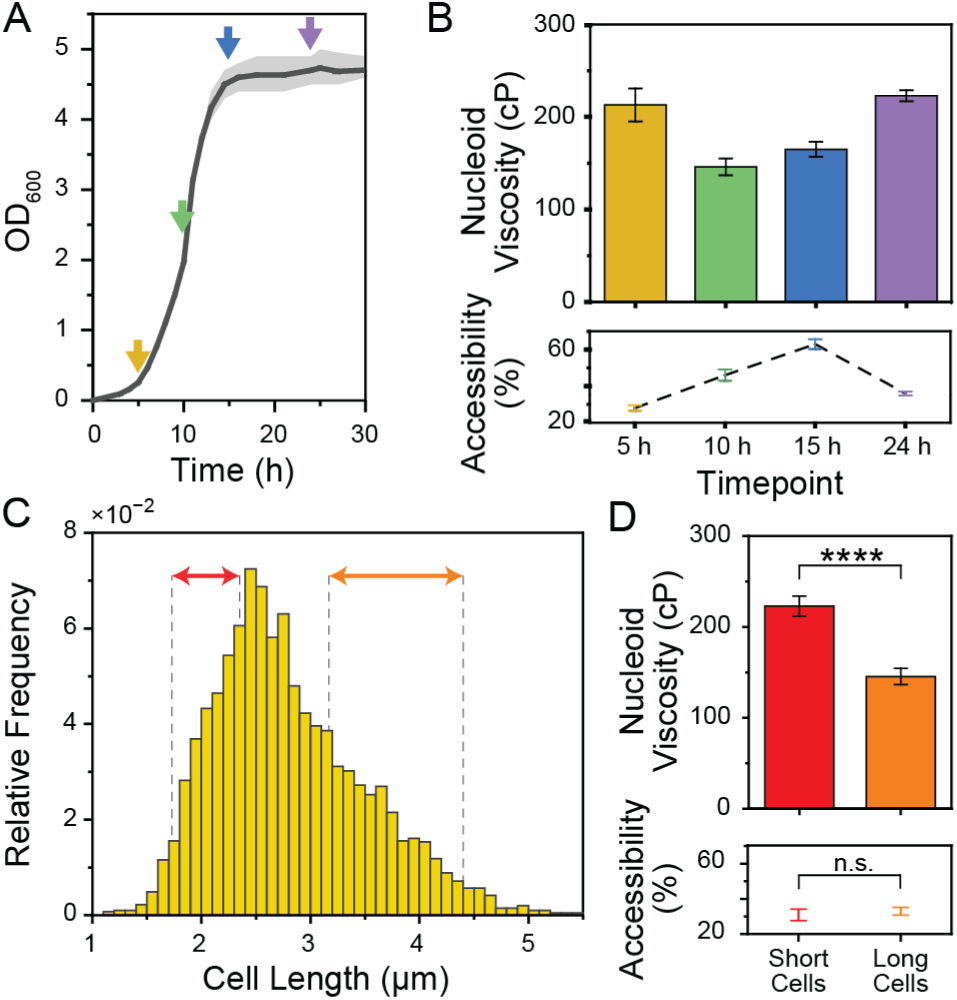
The apparent nucleoid accessibility and effective viscosity are highly correlated with growth phase and cell length. **A**, *E. coli* growth curve. Arrows indicate the time points examined in **B**. Shading: standard deviation from three biological replicates. **B**, Effective nucleoid viscosity and apparent accessibility as a function of growth phase. Error bars: 95% confidence interval (CI) from 50 iterations of bootstrap sampling. Statistical test results are shown in Fig. S3A,B. **C**, Cell length distribution of exponential phase *E. coli* (5 h). Cells with lengths in the 2.5 – 27.5 percentile and in the 72.5 – 97.5 percentile (red and orange arrows, respectively) were analyzed separately in **D**. **D**, Effective nucleoid viscosity and apparent accessibility for the two cell-length distributions for exponential-phase *E. coli* cells (5 h), as indicated in **C**. Error bars: 95% CI from 50 bootstrap samples. Two-sample t-tests with Welch correction between the 2 cell-length distribution groups were performed for nucleoid viscosity and accessibility (n.s.: *p* > 0.05; ****: *p* ≤ 0.0001).

The effective nucleoid viscosity decreases from 5 to 10 h and then increases from mid-exponential phase (10 h) to stationary phase (24 h) (Figs. 2B, S3B): we measured the lowest viscosity during the fastest growth stage, when efficient macromolecular diffusion is necessary for target search within the nucleoid^63,64^. In contrast, prolonged culture and nutrient depletion reduce ATP levels and increase macromolecular crowding^65,66^, which is known to slow protein diffusion^10^, consistent with an increase in effective nucleoid viscosity from 10 to 24 h (Fig. 2B). Interestingly, the effective nucleoid viscosity and apparent accessibility are similar at 5 and 24 h (Fig. 2B). Lag-phase cells retain a stationary-phase transcriptome and metabolic state^67,68^. Consequently, the nucleoid accessibility and viscosity are expected to match the stationary-phase values at the early exponential phase (just as cells exit the lag phase) and then start changing as cells reprogram toward growth. The corresponding apparent cytoplasmic viscosities are presented in Table S9.

Previous studies have shown that DNA replication initiates at a characteristic cell size, such that there is a correlation between cell-cycle stage and cell length during exponential growth^69,70^. We therefore analyzed short cells (1.73 – 2.35 µm, corresponding to the 2.5 – 27.5 percentile) and long cells (3.17 – 4.41 µm, corresponding to the 72.5 – 97.5 percentile) separately, corresponding to populations enriched for early-cell-cycle newborn cells and late-cycle cells approaching cell division, respectively (Fig. 2C). The apparent nucleoid accessibility is similar for both groups (31% and 33%, respectively), but longer cells have significantly less viscous nucleoids than shorter cells (144±9 cP vs. 223±11 cP, *p* < 0.0001; Fig. 2D). We attribute this difference to the nucleoid becoming less constrained and the DNA meshwork becoming more dynamic as the cell elongates and the chromosomes segregate^71,72^, resulting in a more relaxed nucleoid structure with lower effective viscosity as cells approach division.

Across changes in both growth phase and cell cycle, our measurements indicate that nucleoid accessibility and viscosity are properties that vary with cellular state. These findings highlight strong connections between chromosomal physical properties and bacterial growth and division.

### Growth-phase-dependent effects of transcription and translation arrest on the nucleoid properties

To understand how essential biological processes involving the genome influence nucleoid and cytoplasm properties, we inhibited transcription with rifampicin (Rif) and translation with chloramphenicol (Cam) in *E. coli* at different growth phases (Fig. 3). Rif binds to the *β* subunit of RNA polymerase to block transcription elongation^73^ and decreases the mRNA pool in bacteria^74^. Cam binds to the 50S ribosomal subunit to halt polypeptide chain elongation, leaving the ribosome intact^75,76^. We first quantified the properties of exponential phase cells (5 h after subculture, OD_600_ ∼ 0.3; Fig. 2A) following 20 minutes of treatment and found that transcription inhibition by Rif reduces both nucleoid and cytoplasm effective viscosities with a larger effect on nucleoid viscosity (Fig. 3A), while translation inhibition by Cam does not change effective nucleoid viscosity (Fig. 3B). The strong decrease in nucleoid viscosity following Rif treatment is consistent with Rif dissociating polysomes into 30S/50S subunits that can interpenetrate the nucleoid^19,77^, which makes the cytoplasm a better solvent for the chromosome. The decreased cytoplasmic viscosity, which is consistent with previous observations with different-sized probes,^78^ may be due to the decrease in mRNA content that is induced by Rif treatment^74^. Cam treatment stabilizes the polysomes and reduces their disassembly; thus, the polysomes are excluded from the chromosome^19,77^, and we expect nucleoid viscosity to be maintained. Correspondingly, Rif treatment increases the apparent nucleoid accessibility while Cam treatment decreases it (Fig. 3A, B), because of enhanced or weakened DNA-cytoplasm mixing, respectively. Our accessibility measurements are consistent with previous studies showing nucleoid expansion after transcriptional arrest and compaction after translational arrest^18,77^.

**Figure 3.**
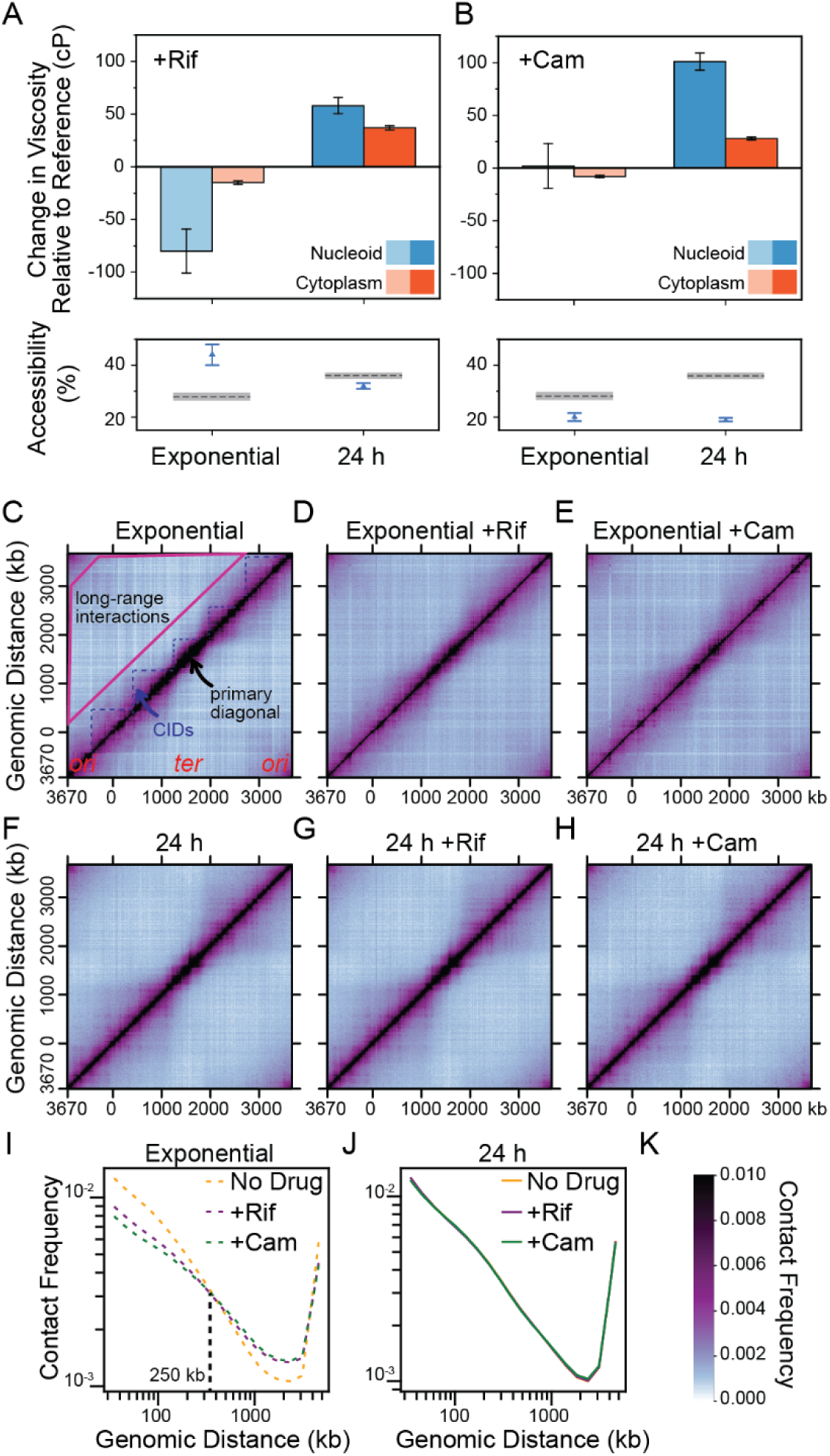
The response of nucleoid viscosity to arresting translation and transcription depends on the growth phases and can be regulated independently of genome organization. **A**, Effects of transcription inhibition by rifampicin (Rif) on the nucleoid and cytoplasm properties at exponential and stationary phases. Top: the effective viscosities after treatment are compared to those in untreated cells at the same growth stage. Error bars: 95% CI derived from bootstrap sampling (Methods). Bottom: the average apparent nucleoid accessibilities of untreated cells (dashed lines) and after treatment (blue triangles). Dashed lines and grey shaded areas: mean and 95% CI for the control conditions. The separate apparent viscosities for the nucleoid and cytoplasm are indicated in Table S10. **B**, As in **A**, but following translation inhibition by chloramphenicol (Cam). **C**, Normalized Hi-C contact maps in 10-kb bins for *E. coli* during the exponential phase without treatment. **D**, as in **C**, but following 30 min of Rif treatment. **E**, as in **C**, but following 30 min of Cam treatment. **F**, Normalized Hi-C contact maps in 10-kb bins for *E. coli* during the stationary phase (24 h) without treatment. **G**, as in **F**, but following 30 min of Rif treatment. **H**, as in **F**, but following 30 min of Cam treatment. **I**, Hi-C contact probability (*P_c_*) curves for cells at exponential phase. The *x*-axis indicates the genomic distance, while the *y*-axis shows the averaged contact frequency. Data are presented at a 10-kb resolution. **J**, as in **I**, but for cells at stationary phase (24 h). **K**, Colorscale bar indicating the normalized contact frequencies for all Hi-C matrices in this study.

Next, we measured cells at stationary phase (24 h). We found that the effective cytoplasm and nucleoid viscosities both increase following transcription and translation inhibition (Fig. 3A,B). This growth-phase-dependent response is likely due to changes in the biophysical environment, the cell metabolism, and the proteomic composition^10,66,79,80^. During exponential phase, the cytosol is actively mixed; arresting transcription or translation in exponential phase can therefore reduce the assembly and abundance of large macromolecular complexes^76,81^, which enables faster probe diffusion. At stationary phase, the lower metabolic level produces a glassy cytosolic environment, and inhibiting gene expression by Rif or Cam treatment further solidifies it^10,82^. Given that the nucleoid is a chromosome body immersed in the cytosol, the changed cytosolic environment contributes to the growth-phase-dependent response of nucleoid viscosity.

Though they cannot resolve properties within specific subcellular regions, we found that ensemble MSD analysis and trajectory-derived diffusion coefficients capture whole-cell trends consistent with our effective-viscosity measurements (Fig. S4). The spatially resolved responses may reflect changes in macromolecular composition and chromosomal structure, as well as changes in energy-consuming cellular processes. Our measurements do not resolve the relative contributions of thermodynamic effects and nonequilibrium activity.

### The nucleoid viscosity does not necessarily vary with changes in genome organization

To understand whether the changes in nucleoid physical properties are correlated with the overall organization and folding of the chromosome, we analyzed genome-wide DNA-DNA interactions by Hi-C for cells grown in the same conditions (Fig. 3C-H).^83,84^ The Hi-C contact map measures the short-range and long-range DNA interactions throughout the genome^85^. All the Hi-C experiments were performed in two biological replicates, which showed reproducible results (Fig. S5). For exponential-phase cells, the Hi-C contact map shows three types of interactions: 1) a strong primary diagonal indicating frequent short-range interactions (Fig. 3C, back pixels on the primary diagonal); 2) mid-range DNA contacts away from the primary diagonal, appearing as squares on the Hi-C contact map and corresponding to chromosome interaction domains (CIDs)^86^ (Fig. 3C, blue dotted lines); and 3) weak long-range interactions further away from the primary diagonal, corresponding to random bumping between distal chromosome loci (Fig. 3C, pink trapezoid). These results are highly consistent with the previously published *E. coli* Hi-C results using different growth media^46,87^.

When cells were treated with Rif, the Hi-C contact map showed an obvious decrease in short-range interactions on the primary diagonal, along with more subtle changes, including reduced CID boundaries and increased long-range interactions (Fig. 3D). This trend is similar to the previously published result of Rif-treated *Caulobacter crescentus*^86^. When cells were treated with Cam, the Hi-C contact map was very similar to that of Rif-treated cells (Fig. 3E). To quantify DNA interactions throughout the genome, we analyzed the average contact frequency between all pairs of loci on the chromosome separated by set distances and plotted the contact probability curve, *P_c_*. Consistent with the Hi-C contact maps, the *P_c_*curves showed similar effects between Rif- and Cam-treated cells: both treatments reduce interactions for loci that are separated by 250 kb or less but increase interactions for loci separated by 250 kb or more (Fig. 3I). The Hi-C results show that, at exponential phase, the inhibition of transcription and translation has similar effects on the folding of the chromosome, leading to reduced short-range interactions and increased long-range interactions. Since active transcription continuously drives DNA supercoiling within the nucleoid^88,89^ and creates CIDs^90–92^, our data are consistent with the idea that halting transcription reduces the local torsional tension (i.e., short-range interactions and CIDs) but increases random bumping of long-range DNA loci. This relaxation of the nucleoid upon transcription inhibition is consistent with our observation of decreased nucleoid viscosity upon Rif treatment at the exponential phase (Fig. 3A). However, for Cam-treated cells, the viscosity of the nucleoid does not change (Fig. 3B) despite the change in chromosome organization (Fig. 3I). These results show that the nucleoid viscosity can remain consistent even when the kilobase-scale genome organization is changed, suggesting that DNA organization is not the sole factor determining the physical property of the nucleoid.

In stationary phase (24 h), transcription and translation levels are already low, and we observed little effect of Rif or Cam on the Hi-C maps and the *P_c_* curves (Figs. 3F-H, J, S5), suggesting that these drugs do not change the overall chromosome organization. Nevertheless, significant viscosity changes are observed in the stationary-phase nucleoid after these treatments (Fig. 3B). Thus, the nucleoid viscosity does not necessarily vary with kilobase-scale genome organization, which raises the possibility that modulation of nucleoid viscosity provides a physical layer to regulate cellular processes, even in the absence of detectable structural reorganization.

### Transcription, translation, and membrane rigidity influence the spatial heterogeneity of nucleoid viscosity

We refined our measurement resolution to the sub-nucleoid scale by targeted masking. The *E. coli* chromosome is hierarchically organized into macrodomains^87,93^ that reproducibly occupy characteristic nucleoid regions^94–97^. We located the most likely positions of the origin (Ori) and terminus (Ter) macrodomains by inserting a *parS* sequence at the target genome region and imaging fluorescent ParB-YGFP, which forms a bright focus nucleated at the *parS*-labeled locus^95,97–99^ (Fig. 4A). From the resulting localization heatmaps, which are widely used to quantify locus distributions^96,100,101^, we identified the top-probability pixels as those comprising 10% of the cumulative probability for each domain (red dashed lines in Fig. 4B). We focused on the spatial distributions of the macrodomains in stationary phase (24 h), when replication initiation is strongly restrained, and the chromosome is less dynamic^102,103,46^, to minimize the domain span and mask overlap. The Ori mask has little overlap with the polar regions where we found the largest discrepancies between experimental and simulated heatmaps. The resulting average positions and local viscosities are shown in Fig. 4B,C. The effective nucleoid viscosity is slightly higher near Ter than near Ori, but overall the two values are very similar (Fig. 4C).

**Figure 4.**
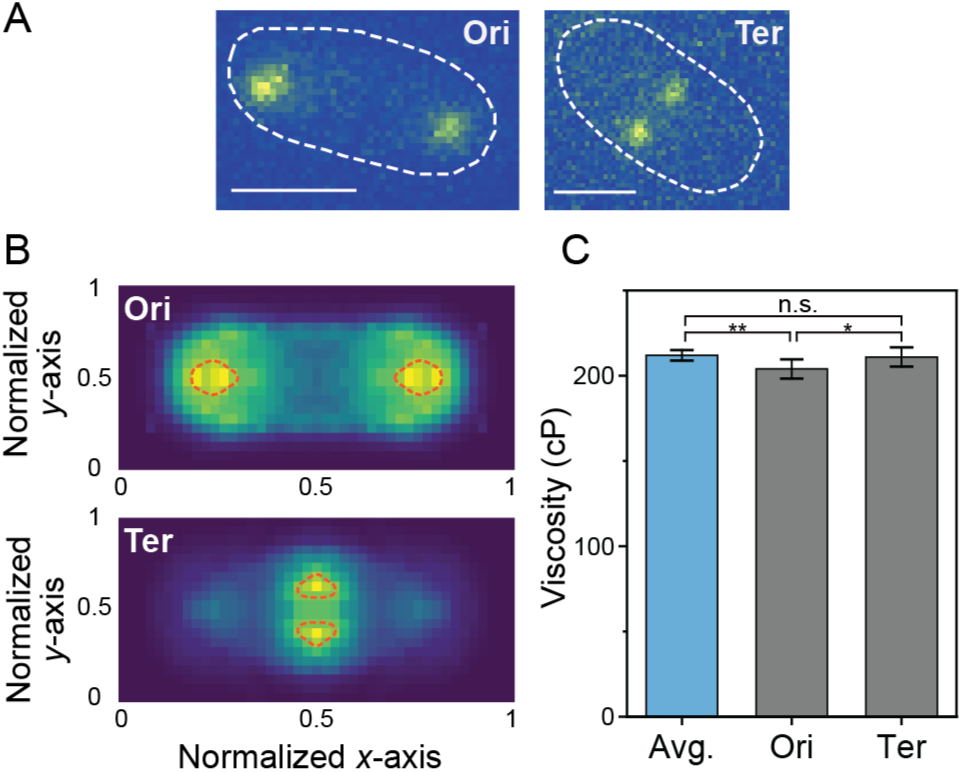
Effective nucleoid macrodomain viscosities at stationary phase. **A**, Example fluorescence images of the origin (Ori) and terminus (Ter) macrodomains labeled by ParB-YGFP at a *parS* sequence. White dashed lines: cell boundaries from the segmentation of the phase-contrast images. Scale bars: 1 μm. **B**, Localization heatmaps of ParB-YGFP foci. Red dashed lines: masks for most likely Ori and Ter macrodomain positions, respectively (Methods). **C**, Average effective viscosity of the whole nucleoid compared to the effective viscosities at the Ori and Ter macrodomains. Error bars: 95% CI derived from 100 iterations of bootstrap sampling (Methods). Comparisons were performed using one-way repeated-measures ANOVA followed by a Tukey post hoc test (n.s.: *p* > 0.05; *: *p* ≤ 0.05; **: *p* ≤ 0.01).

Though we did not detect spatial heterogeneity when comparing the regional effective viscosities surrounding the most probable Ori and Ter locations, prior reports of the relocation of some gene loci toward the nucleoid periphery^104^ and changes in ribosome distribution during gene expression^19,105^ motivated us to partition the nucleoid into periphery and core regions based on geometry to further test for spatial variations in nucleoid viscosity. We defined the nucleoid boundary contour line from the displacement map (Fig. 1B), and we added a margin of 5 – 10% to the nucleoid contour line percentage to generate an inner contour line; the region within this inner contour is referred as the nucleoid core, and the area between the nucleoid boundary and inner boundary is thus the nucleoid periphery. We measured the viscosity of the core and periphery regions in stationary phase (Figs. 5A, S3C), and we found that the core is less viscous than the periphery, with a markedly larger viscosity contrast than that between macrodomains. As the periphery was expanded and the core shrank by increasing the margin from 5 to 10%, the measured periphery viscosity decreased monotonically, indicating a radial viscosity gradient in stationary-phase nucleoids.

**Figure 5.**
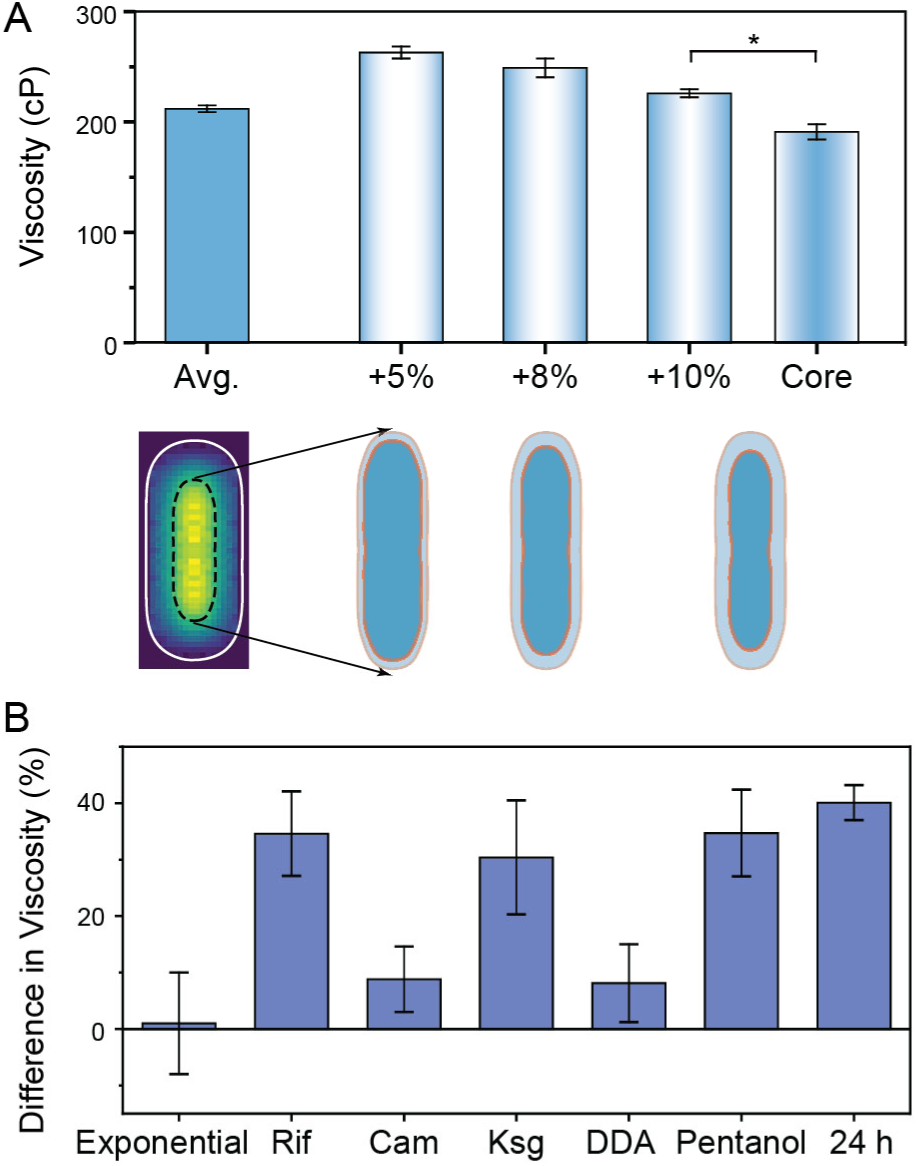
Effects of transcription, translation, and membrane rigidity on the spatial heterogeneity of the nucleoid viscosity. **A**, Effective nucleoid periphery and nucleoid core viscosities at stationary phase measured separately based on defining the core and periphery regions (shown below; dark blue and light blue, respectively), which was realized by adding margin percentages of 5%, 8%, and 10% to the contour line within the displacement map (black dashed line). The nucleoid core effective viscosity is shown alongside the effective nucleoid periphery viscosity for the 10% margin case. Error bars: 95% CI from 50 iterations of bootstrap sampling. The viscosity difference between the nucleoid periphery and core, defined using a 10% periphery margin (Δ*η* = *η*_periphery,10%_ − *η*_core,10%_), was evaluated using a one-sided, null-centered bootstrap hypothesis test (*: *p* =0.031). Statistical tests comparing the whole-nucleoid average and peripheries defined by different margins are shown in Fig. S3C. **B**, Effect of transertion on the effective periphery/core viscosity heterogeneity for the 5% margin case. The percent difference in viscosity, (*η*_periphery_ − *η*_core_)/*η*_whole_ × 100%, was measured at exponential phase and stationary phase, and after Rif, Cam, Ksg, DDA, and 1-pentanol treatment of exponential-phase cells. Error bars: 95% CI from 50 iterations of bootstrap sampling. Statistical test results are shown in Fig. S3D. The separate effective viscosities of the nucleoid core, nucleoid periphery, and cytoplasm are indicated in Table S11.

We hypothesized that this viscosity gradient was related to the coupling of key biological processes. In bacteria, transcription and translation are often physically coupled: ribosomes load onto nascent transcripts and translate them while transcription is still in progress. Furthermore, for genes encoding inner-membrane proteins, the ribosome/nascent-chain complex is recruited to translocons in the inner membrane, enabling co-translational insertion. The concurrent transcription, translation, and membrane insertion is referred to as transertion, which is proposed to link the nucleoid periphery to ribosomes and the inner membrane^106–111^, and we hypothesized that this coupling causes a tension that contributes to the difference between periphery viscosity, *η*_periphery_, and core viscosity, *η*_core_.

To test this hypothesis, we quantified the normalized periphery-core viscosity difference, (*η*_periphery_ − *η*_core_)/*η*_whole_ at exponential phase and stationary phase (Figs. 5B, S3D). At exponential phase, where transertion is more prominent, we found that the difference in viscosities between the nucleoid periphery and core approached 0 at the population scale. Thus, we perturbed transcription, translation, and inner-membrane mechanics in exponential phase cells (Figs. 5B, S3D) to study how breaking transertion-induced tethering influences the heterogeneity of nucleoid viscosity. Rif, which blocks transcription initiation, increased this difference, consistent with the loss of active RNAP-ribosome assemblies and consequent weakening of DNA-membrane coupling at the nucleoid periphery upon rapid transcription arrest^19,20,73^. We inhibited translation with Cam and with kasugamycin (Ksg). Cam does not significantly change the periphery-core viscosity difference, which is consistent with Cam preserving transertion-based tethers, because it stalls elongating ribosomes on existing mRNAs^77,112^ without disrupting the ribosome-DNA connection. By contrast, Ksg increases the periphery-core difference, which is consistent with Ksg binding to 30S ribosomal subunits and interfering with proper mRNA interaction during translation initiation^113^, thereby preventing the formation of ribosome-mRNA complexes and progressively weakening transcription-mediated tethers.

Furthermore, increasing the membrane rigidity with dodecylamine (DDA)^114^ did not change the viscosity gradient, whereas 1-pentanol, which softens the membrane^114,115^, increased the viscosity difference. This viscosity increase by 1-pentanol may be attributed to the higher fluidity of the softened membrane, which increases the lateral mobility of membrane proteins, including translocons^116–118^ and disrupts the transertion-based linkage, in contrast to no disruption to the inner-membrane proteins with ribosome/nascent-chain complexes when the membrane is rigidified by DDA. Furthermore, we compared the exponential phase cells to cells in the stationary phase (24 h), when global transcription/translation and membrane insertion are reduced^61,119,120^. The contrast between periphery and core viscosities is greater in stationary phase than in exponential phase without perturbation (Figs. 5B, S3D). The chemical perturbations used here can affect multiple cellular processes. For example, 1-pentanol can also influence transcription, translation, and cytoplasm organization. However, we found that conditions expected to break the tethering (Rif, Ksg, 1-pentanol, and stationary phase) amplify the periphery-core viscosity difference, whereas conditions that minimally disrupt linkages (Cam, DDA) do not significantly change the periphery-core viscosity difference relative to the untreated control. These data are compatible with a model in which a reduction of linkages between RNAP, the ribosomes, and the inner membrane in stationary phase contributes to a larger periphery-core viscosity difference, whereas stronger coupling in exponential phase is associated with more uniform viscosity.

## Discussion

### A readily implementable and generalizable framework for quantifying the biophysical properties of the bacterial nucleoid

This work introduces a quantitative, generalizable microrheological framework for determining the biophysical properties of the bacterial nucleoid by combining 2D single-particle tracking with 3D BD simulations to characterize the apparent nucleoid accessibility and effective viscosity. Although conventional 2D imaging setups are widely used for intracellular microrheology^6,8,10,12,26,121^, these microscopes collapse 3D positions onto a 2D plane. This projection obscures the influence of the complex cell geometry and can lead to biased estimates of dynamics and localization distribution^122,123^. Valverde-Mendez et al. used 3D imaging to show that geometric confinement strongly affects the diffusion of ribosome-sized particles and uncovered probe distributions that are not apparent in 2D projections^31^, underscoring that diffusion in the true 3D cellular volume cannot be neglected. However, applying full 3D imaging to quantify nucleoid viscosity at the population scale and across many conditions is experimentally demanding and typically requires specialized instrumentation^124^, limiting its routine use in many laboratories. Our framework addresses this gap by using a standard epifluorescence microscope to record 2D trajectories and then employing 3D BD simulations to reconstruct the underlying 3D probe motion statistically, thereby differentiating motion within the nucleoid from cytoplasmic contributions to the measured diffusion. We use the Stokes-Einstein relation operationally to convert the measured probe diffusivity into an effective viscosity that quantitatively characterizes the local material environment experienced by the probe. Although previous studies have reported deviations from classical Stokes-Einstein scaling for protein diffusion in the bacterial cytoplasm^47^, our use of the same probe, imaging conditions, and analysis provides a consistent basis for comparing effective viscosity estimates across regions and perturbations.

This framework is compatible with other genetically encoded passive microrheology probes, such as GEM particles^8^, and can be extended to mesoscale probes engineered by artificial protein design^125,126^. Because it does not rely on *E. coli*-specific biology and only requires a defined cellular morphology, it can, in principle, be applied to other bacteria with known shapes. The modest optical requirements and high throughput of this approach make quantitative measurements of bacterial chromosomal biophysical properties broadly accessible and enable systematic studies of the coupling of these properties to cellular processes and physiology.

### What factors regulate the material properties of the bacterial nucleoid?

Active adaptation of the cytoplasmic viscosity has been demonstrated: yeast cells modulate their cytosolic viscosity by changing glycogen and trehalose levels^127^, mTORC1 signaling can tune the ribosome concentration, and thus molecular crowding^8^, and in *E. coli*, the cytoplasmic viscosity is strongly correlated with the abundance of amino acid metabolism proteins^12^. Our results show that the nucleoid viscosity and accessibility change as a function of growth phases and the cell cycle. This regulation of the nucleoid material properties is consistent with the changing functional demands of the cell under different conditions.

The nucleoid can be viewed as a phase-separated, active viscoelastic hydrogel. A related hydrogel framework has been proposed for chromosome organization in human nuclei, where an RNA-protein mesh forms interconnected microgels that influence chromatin compaction and protein mobility^128,129^. Rheology theory predicts that its viscosity is primarily governed by the DNA concentration^130–132^ and the degree of cross-linking^130,133^. Changes in the nucleoid volume can substantially shift its viscoelastic state by altering the effective DNA volume fraction, and NAPs could tune the nucleoid viscosity by modulating the cross-linking density. In this hydrogel picture, the nucleoid is a meshwork immersed in a crowded liquid medium; macromolecular crowders generate depletion forces that induce short-range attractions between DNA segments, thereby increasing internal friction, which could influence the bulk viscosity of the nucleoid^134,135^.

Beyond the hydrogel picture, the nucleoid is an active system that is persistently remodeled by proteins like SMC complexes, topoisomerases, and RNA polymerase^13,136^. The *E. coli* MukBEF complex can reel DNA into loops, making a bottle-brush-like chromosome architecture^137^ that tends to reduce inter-chain entanglement and friction^138^. Topoisomerases regulate the level and spatial distribution of supercoiling^139^, which changes the entanglement in the plectonemic chromosome^140,141^. In addition to cross-linking DNA, some NAPs can alter DNA persistence length^142^. Polymer physics suggests that these properties can contribute to changes in nucleoid viscosity.

Therefore, we hypothesize that, in contrast to the cytoplasm—which is often described as liquid-like near the glass transition^10^ and can regulate viscosity by adjusting the concentration of macromolecules^12,127^—the nucleoid can adapt its material properties by regulating NAP expression and recruiting MukBEF complexes and topoisomerases to change the DNA compaction, cross-linking density, and DNA conformation. Our measurements of the effective nucleoid viscosity across growth phases, cell-cycle stages, and under transcriptional perturbations are consistent with this model. This coarse-grained adjustment of the viscosity enables the nucleoid to maintain the chromosome shape against turgor pressure while ensuring that the environment is sufficiently fluidized for essential processes like gene expression and DNA segregation.

### The interplay between cellular processes and the biophysical properties of the nucleoid

Our results further show that the biophysical properties of the nucleoid respond to transcription and translation inhibition. Moreover, inhibiting transcription or translation produces opposite changes in the effective nucleoid viscosity in exponential vs. stationary phase cells, which supports our model that the regulation of nucleoid viscosity depends sensitively on the underlying biomolecular composition, crowding, and spatial organization. Though drugs work on the same molecular targets in all growth phases, the biomolecular environment is distinct, and thus, the nucleoid viscosity response to drugs is strikingly different. Additionally, by comparing these biophysical measurements to Hi-C assays, we found that nucleoid viscosity can be adjusted without detectable changes in the kilobase-scale DNA-DNA interactions, which highlights that material properties constitute an independent regulatory axis.

Upon refining our characterization to the sub-nucleoid scale, we also found a pronounced spatial heterogeneity in the effective nucleoid viscosity. Differences between macrodomains are much less pronounced than the contrast between the periphery and the core, implying that the local viscosity is more likely to be regulated according to the spatial position within the nucleoid rather than by the underlying DNA sequence. Though it has not been directly visualized^111^, transertion is consistent with our observations: changing the inner membrane rigidity affects the periphery-core viscosity contrast in the same way as breaking tethers at the nucleoid or ribosomes. This similarity indicates a direct link from the chromosome, through ribosome-mRNA complexes, and to the inner membrane. Moreover, disrupting these linkages in exponential-phase cells produces a similar periphery-core viscosity contrast as was measured in stationary-phase cells, which suggests that transertion plays an important role in the spatial regulation of nucleoid viscosity.

## Limitations of the study

This study characterized the biophysical properties of the nucleoid in *E. coli* cells grown with glycerol as the carbon source. The reported quantities are apparent accessibilities and effective viscosities experienced at the 25-nm-probe scale, rather than probe-size-independent properties of the nucleoid. Generalization across length scales would require systematic measurements using probes of different sizes. These quantities represent how the chromosomal environment influences dynamics of biomacromolecules at this relevant scale. The possible time-scale dependence associated with anomalous transport is not resolved by the present analysis. In our current framework, the effective viscosities are inferred by converting probe displacements over the experimental sampling interval to viscosities with the Stokes-Einstein equation. Thus, the effective viscosity may reflect effects of confinement, transient interactions, or subdiffusive dynamics. The present analysis does not distinguish these contributions or resolve possible time-scale dependence associated with anomalous transport.

Our framework should apply readily to other bacteria with well-defined geometries. We anticipate that future studies will indicate differences in the physical properties of the nucleoid in *E. coli* in other nutrient regimes and in other model organisms. Furthermore, single-cell measurements will extend the current results beyond population-level statistics and directly measure cell-to-cell heterogeneity. This extension will require a greater number of tracks per cell for robust inference. Additionally, our BD simulations capture probe diffusion, but could be improved by incorporating spatial gradients in cytoplasmic crowding and by simulating an ensemble of cells with a distribution of shapes drawn from the experimentally measured morphology. Finally, we consistently observed a higher effective viscosity at the nucleoid periphery than at the core in stationary phase and upon transertion inhibition; the mechanism underlying this spatial pattern is not yet understood.

## Conclusions and future perspectives

This work demonstrates a paradigm for investigating biological processes in the context of their physical and material environment in living cells. We find that the bacterial nucleoid is embedded in a complex, dynamic material environment, and our results highlight how both local and global nucleoid mechanics are coupled to essential processes such as gene expression, growth, and stress adaptation. This perspective adds a physical layer to the regulatory landscape of bacterial chromosome organization, which complements traditional models based on sequence, structure, and biochemistry.

Quantitatively measuring the nucleoid physical properties further provides essential constraints for synthetic-cell design and computational modeling^43–45^. Reproducing life-like processes in artificial matrices will require coordinated regulation of not only biochemical pathways, but also the physical properties of the environment^143,144^. Absolute measurements of viscosity and accessibility supply critical parameters for simulations and theoretical models, enabling more realistic descriptions of reaction-diffusion dynamics and chromosome mechanics^145–147^. The bacterial nucleoid is a micron-scale, phase-separated condensate, which is comparable to chromatin condensates in eukaryotic systems^148,149^. Thus, dissecting its rheological properties and their molecular control is likely to yield insights that generalize to genome-associated condensates more broadly, thereby contributing to the growing field of phase separation and material states within the genome.

Ultimately, our results demonstrate that the biophysical properties of the nucleoid differ markedly from those of the surrounding cytoplasm, underscoring the importance of treating the nucleoid as a distinct physical compartment. By quantifying the apparent nucleoid accessibility and effective viscosity (1) across growth phases and cell-cycle states, (2) under transcription and translation inhibition in both exponential and stationary phases, and (3) upon perturbation of different components of transertion-mediated linkages, we showed that the bacterial nucleoid is dynamically remodeled, spatially heterogeneous, and highly responsive to perturbations of cellular processes.

## Materials and Methods

### *E. coli* Cell Growth

All strains used in this study were derived from prototrophic *E. coli* K-12 strain W3110 (CGSC no. 4474). Cells were streaked from glycerol stocks onto LB-agar plates containing antibiotics at the following concentrations: 20 μg/mL kanamycin (CAS no. 25389-94-0) for the strains with fluorescently labeled ParB*-parS* DNA loci or 100 μg/mL spectinomycin (CAS no. 22189-32-8) for the nanocage-GFP strain. Single colonies were isolated from plates and grown in Luria-Bertani (LB) Broth (Fisher BioReagents) supplemented with corresponding antibiotics and incubated at 37°C overnight. The liquid culture was then back-diluted to a factor of 1:100 into High-Def Azure (HDA) medium (Teknova cat. No. 3H5000) supplemented with 0.2% glycerol (m/v) and incubated at 30°C. All shaking was at 250 rpm. The strains, plasmids, oligonucleotides, and Next-Generation sequencing are listed in the SI Appendix, Tables S1 – S4.

### Fluorescence Microscopy

#### Single-particle tracking of sfGFP-labeled nanocages

sfGFP-labeled protein nanocages were expressed from an arabinose-inducible promoter. For the 5-h and 10-h time points, the back-diluted culture was grown to OD 0.3 at 37°C, then arabinose was added to a concentration of 0.007% – 0.010% or 0.030% – 0.040%, respectively. When the OD reached 2, the culture was diluted 100× a second time in HDA medium and incubated at 30°C for an additional 5 h or 10 h. For the 15-h and 24-h time points, the back-diluted culture was grown at 30°C to OD 0.15 – 0.20, then arabinose was added to a concentration of 0.0008% – 0.0010% or 0.0012% – 0.0016%, respectively. We adjusted the arabinose concentration within a small range to maintain a sparse probe density suitable for reliable single-particle localization and tracking across different experiments. A single fixed concentration can produce too few detectable probes in some cultures or excessively dense, overlapping fluorescence signals in others due to variations in nanocage expression between batch cultures and growth phases. We therefore used this narrow range of arabinose concentrations to compensate for culture-to-culture variation and ensure consistent, high-fidelity probe tracking under all conditions.

After the specified incubation time, spent medium was prepared by centrifuging 3 mL of saturated culture from each culture tube for 4 min at 7200 × g, then syringe filtering the supernatant at least twice using a fresh 0.22-μm syringe filter (Fisherbrand) each time. Then, cells were concentrated 10 – 100×, and 2 μL of concentrated cells were pipetted directly onto a 2% (w/v) agarose pad made from spent medium and the specified antibiotics or chemical (Table S5) and covered by a No. 1 coverslip (Corning).

Imaging was performed on an Olympus IX-71 inverted microscope with wide-field epifluorescence illumination with a 100× 1.40 NA oil-immersion objective heated to 30 °C by an objective heater (Bioptics) and using appropriate index-matched oil (Zeiss). The cell sample was mounted on the microscope objective and left to rest for 20 min to reach thermal equilibrium and permit drug/chemical treatment as specified before imaging. Each sample was imaged for 60 – 85 min. Fluorescence imaging of the sfGFP-labeled nanocages was performed with a 488-nm laser (Coherent Sapphire 488-550) and a dual-band filter (Chroma, ZET488/561 M-TIRF) with a power density of 60 – 90 W/cm^2^.

Based on the axial Rayleigh formula for the depth of field, *DOF* = 2*n*/*NA*^2^, for our high-numerical-aperture objective (NA = 1.49), refractive index-matching oil, and emission wavelength of ∼500 nm, the minimum depth of field extends ±0.34 µm from the focal plane. Centering the focal plane at the mid-plane of the cell provides a baseline in-focus range of 0.68 *μ*m. Because our 25-nm nanocage probes emit high photon counts relative to background noise, our single-particle tracking algorithm consistently detects valid probe localizations even when axial displacement introduces moderate out-of-focus point spread function broadening. By applying an empirical point spread function width cutoff greater than the diffraction limit during particle tracking, we consistently track nanocage probes across the entire 1.0-µm cell diameter.

Movies capturing nanocage diffusion were acquired continuously with a 512 × 512-pixel Photometrics Evolve EMCCD camera (40-ms frame time).

#### Chromosomal locus imaging

Cells expressing ParB-YGFP and containing the *parS* sequence at specific chromosomal loci were incubated at 30°C in HDA medium after back dilution. After 24 h, concentrated cell samples were mounted on agarose pads made from spent medium as described above for nanocage tracking. Fluorescence microscopy of the ParB-YGFP was performed as described above for nanocage tracking, with excitation from a 488-nm laser (Coherent-Sapphire 488-50) and dual-band filter (Chroma, ZET488/561 M-TIRF) at a power density of 180 W/cm^2^ and appropriate optical filters.

### Hi-C Experiments

Cells used in Hi-C were grown in the same procedure as described for “Single-particle tracking of sfGFP-labeled nanocages”. Detailed procedures and data analysis can be found in the SI Appendix.

## Supplementary Methods

All bacterial strains, plasmids, image processing, data analysis, and simulations used in this study are described in the SI Appendix.

## Acknowledgements

The nanocage plasmids were a generous gift from Christine Jacobs-Wagner. We thank the Indiana University Center for Genomics and Bioinformatics for high-throughput sequencing. Support for this work comes primarily from National Institutes of Health Grant R01 GM144731 to J.S.B. and R01 GM143182 to J.S.B. and X.W. Additional support came from National Institutes of Health R01 GM141242 (X.W.) and R01 AI172822 (X.W.). This research is a contribution of the GEMS Biology Integration Institute, funded by the National Science Foundation DBI Biology Integration Institutes Program, Award #2022049 (X.W.). L.A.M. gratefully acknowledges support from the National Institutes of Health K12GM111725.

## Author contributions

X.D. developed the experimental and analysis framework, designed, performed, and analyzed all live-cell imaging and tracking experiments and simulations, and wrote the first manuscript draft. L.A.M. constructed plasmids and strains. L.E.W., E.E.W., and Q.L. performed and analyzed Hi-C experiments. A.M.J. performed nucleoid-staining experiments and analyzed the data. X.W. and J.S.B. designed experiments and supervised the study. All authors wrote and edited the manuscript.

## Data and code availability

Hi-C data were deposited to the NCBI Gene Expression Omnibus (accession no. GSE314147; reviewer token aduhkwgkhdcbhin). The raw microscopy data reported in this study cannot be deposited in a public repository due to its large size. The full dataset and intermediate outputs for individual conditions are available upon request from the lead contact with no restriction. A subset of the nanocage single-particle tracking data and all locus-tracking data were deposited to Zenodo (https://zenodo.org/records/18599214; https://zenodo.org/records/18598888). Data analysis tools are available on the Biteen Lab GitHub at https://github.com/BiteenMatlab/Nucleoid_Viscosity.

