## Supplemental Information for "The biophysical properties of the bacterial nucleoid are dynamic, heterogeneous, and responsive to perturbations of cellular processes"

**Contents:**

- **Supplementary Methods**
- **SI Figures S1 – S10**
- **SI Tables S1 – S12**
- **Supplementary References**

### Supplementary Methods

#### Image processing, Data analysis, and Simulations

*Cell segmentation*

Cell segmentation was performed on phase-contrast images using the Cellpose[1] package, trained on *E. coli* cells. These segmentation masks were used for tracking and cell morphology analysis in the downstream processes.

*Single-particle localization and tracking*

Detection, localization, and tracking of single fluorescent nanocages and ParB-labeled macrodomain foci were conducted using the SMALL-LABS algorithm[2].

*Probe localization heatmaps and displacement maps*

All 24-h measurements include >4000 cells, and all exponential-phase measurements include >1000 cells. The 10-h measurement included 5077 cells, and the 15-h measurements included 6762 cells. The long cell vs. short cell comparison (Fig 2. C,D) includes 1010 and 1006 cells, respectively. Two biological replicates were combined into one dataset for each condition to attain the required amount of data (>$1.0 \times 10^6$ localizations and >$7.5 \times 10^5$ steps for each condition), as we described previously[3,4]. We used bootstrapping to calculate the biological variance.

For each segmented cell, the length (long axis) and width (short axis) were calculated from the maximum and minimum Feret diameters of the cell mask, respectively. To ensure morphological consistency across the population, a length-based filtering criterion was applied for each dataset unless otherwise noted. Cells were retained if their lengths fell within the range defined by the mean ± the smaller of the standard deviation of the cell length distribution or 0.75 μm in exponential phase and 0.5 μm in stationary phase. The heatmap and displacement map for each condition were generated using the average aspect ratio of cells passing the length filter.

To generate localization heatmaps, the coordinates of fluorescent probes detected within each cell were normalized by the cell's respective long and short axis dimensions. These normalized coordinates were compiled across all cells into a two-dimensional histogram to visualize the ensemble localization density. Though the random cell orientations produce a reasonably symmetric ensemble-averaged map, finite sampling can still result in unequal numbers of observations and residual fluctuations between corresponding spatial bins. The resulting heatmaps were therefore symmetrized along the long and short axes and normalized such that the sum of the localization occurrences is 1. This procedure enforces the reflection symmetry assumed for the ensemble-averaged cell and nucleoid geometries and reduces sampling noise.

Displacement maps were computed using displacements from the tracking data (change in position per 40-ms frame time). Each displacement was associated with a midpoint position, which was normalized using the same coordinate transformation as in the heatmaps. We found minimal differences when assigning each displacement to the starting point instead (Fig. S6). Displacements were then binned into the same 2D grid used for the localization maps. The map was symmetrized along both axes. The mean step size of all displacements falling within each bin was computed and assigned as the bin value. Grid bins containing fewer than 5% of the maximum number of steps were omitted.

The localization and displacement maps were divided into 20 spatial bins across the mean cell width for each condition. The number of bins along the cell length was then determined from the corresponding mean cell aspect ratio. Thus, the spatial bin size was condition-dependent and was calculated as the mean cell width divided by 20. Given a typical *E. coli* cell width of approximately 1 μm, the average bin size was approximately 50 nm.

*Cell boundary and nucleoid outline determination*

To enable accurate 3D reconstruction of cell geometry, smooth outlines of the cell and nucleoid were calculated from the localization heatmaps and displacement maps. The example for untreated, exponential-phase cells is shown in Fig. S7A.

The heatmaps were interpolated using the *interp2* function in MATLAB (2024a) and Gaussian-smoothed (Fig. S7B). In the cases where this interpolation magnified small point-to-point fluctuations into oscillatory artifacts that disturbed the subsequent gradient fitting, the more robust but more computationally expensive kernel density estimation (KDE) approach was used.

The boundary obtained from phase-contrast images represents the overall cell envelope and does not resolve the inner membrane or the volume accessible to the 25-nm probes. The gradient of the probe-localization map was therefore used to identify the sharp decrease in localization density near the cell periphery. This process provides a boundary of the cytoplasmic region actually sampled by the nanocage probes.

Gradients of the smoothed heatmap were computed in the *x* and *y* directions. The gradient magnitude map was then calculated as the square root of the sum of the squared gradients in both directions in each bin (Fig. S7C). The average value of the histogram of the gradient magnitudes in the valley between the peaks was identified (yellow dashed line in Fig. S7D), and all gradient values above this baseline were fit to a Gaussian mixture model (green solid line in Fig. S7D). Only pixels in the gradient map (Fig. S7C) exceeding a first threshold, defined as the mean plus two standard deviations of the lower Gaussian peak (Fig. S7D, red dashed line), are plotted in Fig. S7E.

We used a two-component Gaussian mixture model as an automated image-segmentation tool rather than an inferential model to estimate the physical parameters. The initial thresholds, defined by mean + 2× standard deviation served to establish preliminary candidate regions; we subsequently refined the final nucleoid boundaries using local directional-gradient positions and displacement contours as described below.

The outer boundary of the region in Fig. S7E corresponds to the inner membrane boundary, beyond which no nanocage signal is expected (Fig. S7E, red dashed line). We defined this outermost edge as the cell boundary. Pixels in Fig. S7C within this cell boundary and that have a gradient below a second threshold, defined as the mean minus the standard deviation of the higher Gaussian peak in Fig. S7D (blue dashed line), are plotted in Fig. S7F. This region comprises an outer circle adjacent to the cell boundary and a disconnected inner region (Fig. S7F, region within pink outline); the inner region was used as the starting point for further steps to determine the nucleoid boundary.

To further refine the nucleoid boundary definition, we identified the nucleoid exclusion zone piecewise. Throughout most of the cell, exclusion of cytoplasmic nanocages by the nucleoid occurs in the $y$ direction, so the nucleoid edge was identified as the sharpest gradient in the $y$ direction. The horizontal line profile of occurrences along $y = 0.5$ within the pink boundary (black arrow in Fig. S7G) is plotted in Fig. S7H, and the two steepest positions between the maxima were used to define the $x$-scan range (Fig. S7G, H, gray lines). Vertical line profiles of occurrences within this range were plotted; example line profiles along $x = 0.30$, 0.50, and 0.75 (blue dotted lines in Fig. S7G) are plotted in Fig. S7I. For each $x$ value, the coordinates with the steepest change between occurrence maxima were chosen (Fig. S7I, red crosses), and these points form the $x$-scan components of the exclusion region boundary (brown lines in Fig. S7L). On the other hand, at the nucleoid caps, exclusion occurs in all directions, so the nucleoid boundary was calculated using the gradient value (Fig. S7J). Horizontal line profiles of the gradient were plotted for $y$ values within the gap between the upper and lower brown lines in the boundary lines in Fig. S7M (white lines). For each $y$ value, the maxima in the gradient line profile along the $x$ axis (Fig. S7K, black crosses) are chosen to form the $y$-scan component of the exclusion region boundary (Fig. S7M, brown lines). Finally, the boundary components in Fig. S7L and M were combined to define the full exclusion region (Fig. S7N).

This exclusion zone was further refined to estimate the nucleoid boundary based on information from the displacement map (Fig. S7O). The displacement map was interpolated using the *interp2* function and smoothed with a Gaussian kernel (Fig. S7P). The smallest contour line that fully encloses the exclusion region (brown line in Fig. S7P) was identified (black solid line in Fig. S7P). We note that this contour line is very similar to the largest contour line fully contained within the exclusion region (black dashed line in Fig. S7P) and to the exclusion zone boundary itself (brown line in Fig. S7N, P). The nucleoid outline used for further analysis was defined as the outer contour line.

*3D cell geometry reconstruction*

For each growth phase and treatment, the cell boundary and nucleoid outline were determined from the experimental heatmap and displacement map. In 2 dimensions, the cell boundary was modeled as a rectangle with semicircle ends (Fig. S7Q, cell boundary: red dashed line; fit: blue solid line). The nucleoid boundary was represented as a piecewise linear function, with each segment defined by two adjacent points along the experimental outline (Fig. S7Q, black line). Specifically, we first interpolated the experimental displacement map ten-fold and applied Gaussian smoothing before using contour analysis to define the two-dimensional nucleoid boundary. We then connected adjacent boundary coordinates using straight-line segments to form a piecewise-linear outline of the nucleoid. The cell and nucleoid geometries were then reconstructed in 3D based on their rotational symmetry along the $y$ = 0.5 line (Fig. S7R). The reconstructed 3D cell volume was segmented into two regions: the cytoplasm (between the cell envelope and the nucleoid surface) and the nucleoid (within the nucleoid surface).

*3D Monte Carlo simulations of probe positions and nucleoid accessibility measurement*

We used a Monte Carlo procedure as a static position-sampling method to infer the apparent nucleoid accessibility of the 25-nm probe. To simulate uniform probe distributions, $2 \times 10^6$ probes were randomly placed throughout the entire cell volume, resulting in equal initial densities in the cytoplasmic and nucleoid regions. Specifically, candidate ($x$,$y$,$z$) coordinates were sampled uniformly within a rectangular volume enclosing the reconstructed cell. Candidate positions outside the cell boundary were rejected, and this procedure was repeated until the specified number of probes had been placed throughout the entire cell volume. A series of static simulations was then performed across a range of accessibility values. Nucleoid accessibility was defined as the ratio of the probe density in 3D within the nucleoid to the 3D probe density in the cytoplasm. By this definition, an accessibility of 100% indicates complete probe penetration into the nucleoid with no exclusion, whereas 0% indicates total exclusion of probes from the nucleoid. For each accessibility value, a simulated localization heatmap was generated using $2N_{exp}$ probe positions randomly sampled from the pool of $2 \times 10^6$ simulated positions, where $N_{exp}$ is the number of localizations in the corresponding experimental heatmap. Only the corresponding fraction of the probes within the nucleoid region was randomly retained, while all probes in the cytoplasm were kept. These combined probe sets were used to generate simulated localization heatmaps.

Simulated localization heatmaps for a range of accessibility values (from 0% to 100% in 1% increments) were compared to the experimental heatmap using the Structural Similarity Index Measure (SSIM). The resulting SSIM values were plotted against accessibility, and the accessibility value corresponding to the peak of a fifth-order polynomial fit curve was taken as the most likely nucleoid accessibility for that condition.

*3D Brownian dynamics simulation within the reconstructed cell geometry*

To simulate probe diffusion, particles were initialized at random positions within the cytoplasm. A modified discrete-time Brownian random-walk model, which incorporates confinement via rigid collisions at both the cell membrane and the nucleoid boundaries, was employed. At each step, the displacement magnitude, *r*, was drawn from a Rayleigh distribution about a median set by the parameter *step_size*. The direction was randomly sampled in spherical coordinates: the azimuthal angle, $\phi$, was sampled from a uniform distribution in [0,2π), and the polar angle, $\theta$, was sampled from a uniform distribution within the range [−π,π].

When a proposed step crossed a boundary, a rigid reflection was applied, preserving the original length *r*. This boundary condition ensured that all probe trajectories remained confined within the cytoplasmic region, consistent with experiments. The simulation proceeded until $1.2 \times 10^5$ steps were completed.

*Nucleoid viscosity calculations*

The effective nucleoid viscosity was determined by comparison to BD simulations within the reconstructed 3D cell geometry using a range of average step sizes from 5 – 40 nm in 5-nm increments. For each step size, experimental imaging was simulated by sampling probe positions at 5- to 250-step intervals (in 5-step increments), and simulated displacement maps were generated by retaining the probe positions at each sampling interval.

These simulations recapitulated 3D probe diffusion in the cytoplasm, but diffusion within the nucleoid was not simulated. Therefore, agreement between simulated and experimental displacement maps was assessed using a custom difference score that focuses on cytoplasmic diffusion behavior and spatial confinement by the inner membrane and nucleoid boundaries. The difference score consists of two components:

1. $l^2$-distance of step size differences: For each grid bin within the cytoplasm region, the $l^2$-distance in average displacement between the experimental and simulated displacement maps was calculated. For each included cytoplasmic bin (*i,j*), the weight was calculated as: $w_{i,j} = N_{i,j}/\sum_{(k,l)\in C} N_{k,l}$ , where $N_{i,j}$ is the number of experimental displacement steps assigned to bin (*i,j*), and $C$ denotes all

cytoplasmic bins included in the $l^2$-distance calculation. The weights were therefore normalized to sum to one across the included bins, giving greater influence to bins supported by more displacement observations.

2. Pearson correlation coefficient (PCC): To capture global spatial patterns, the PCC between the experimental and simulated displacement maps was calculated for all bins, and cytoplasmic bins were given a 25% weight boost prior to normalization, biasing the pattern comparison toward the cytoplasmic region.

The final difference score was computed as a weighted sum of the two metrics:

$$S = w_1 d_{L2} + w_2 PCC$$

Where the $l^2$-distance, $d_{L2}$, is weighted with $w_1 = 0.5$, and the PCC is weighted by an empirically chosen weight, $w_2 = -120$ (Fig. S8A-D). The positive sign on $w_1$ and the negative sign on $w_2$ ensure the convention that a smaller difference score indicates better agreement between experimental and simulated displacement maps.

The weights $w_1$ and $w_2$ were chosen to make $w_1 d_{L2}$ and $w_2 PCC$ comparable in magnitude across the explored parameter space, and we tuned $w_2$ relative to $w_1 = 0.5$ to prevent bias on a single component. When $|w_2|$ is too large (e.g., −180 or −240 in Fig. S8A-D), the difference score is PCC-dominated (the minima in difference score vs. sampling interval or step size are much larger than the target range found in the plot of $l^2$-distance vs. sampling interval or step size). Conversely, when $|w_2|$ is too small (e.g., −60 in Fig. S8A-D), the difference score is $l^2$-dominated (minima in difference score vs. sampling interval or step size are much smaller than the target range found in the plot of PCC vs. sampling interval or step size). We therefore selected the intermediate value of $w_2 = -120$, which consistently selected parameter values that simultaneously meet pattern fidelity (high PCC) and displacement agreement (low $l^2$-distance).

The simulated map with the lowest difference score was selected for downstream analysis. The fitting and analysis of displacement distributions from the experimental and simulated data are described in the main text (Equations 1 – 3).

To fit the experimental displacement distribution (Fig. 1I, middle) to a two-component Rayleigh mixture, the weights $w_{\text{cyto}}$ and $w_{\text{nuc}}$ were estimated based on the nucleoid accessibility. Specifically, in the corresponding static simulation with the corresponding optimal accessibility, all probes whose ($x$, $y$) positions fall within the nucleoid mask were projected into the nucleoid region in the 2D map. The 3D position was used to categorize these positions as cytoplasm or nucleoid, and the fraction of the cytoplasmic portion was taken as $w_{\text{cyto}}$. The remaining weight, $w_{\text{nuc}} = (1 - w_{\text{cyto}})$, corresponds to the nucleoid-confined population. To allow for fitting flexibility, the Rayleigh parameter (from simulation) was allowed to vary over a ±15% range, and $w_{cyto}$ was allowed to vary within ±7.5% of the estimated value.

The localization-derived accessibility, together with the reconstructed 3D geometry, was used to determine the cytoplasmic and intra-nucleoid weights for decomposition of the experimental displacement distribution. Considering that the localization and displacement data are derived from the same pool of trajectories, and $2 \times 10^5$ tracked localizations were used to generate each map to minimize sampling uncertainty, the ratio between the number of displacements in different areas matches the ratio between the number of localizations.

*Cytoplasm viscosity calculations*

The probability density function (pdf) of experimental displacements within the cytoplasm region was fit to a Rayleigh distribution (Eq. 1). The resulting fit parameter, $\alpha$, was used to calculate the diffusion coefficient, $D$ (Eq. 2), and the effective cytoplasmic viscosity, $\eta$, was subsequently determined via the Stokes-Einstein equation (Eq. 3), following the same procedure as for nucleoid viscosity estimation.

*Sub-organelle nucleoid viscosity*

The viscosity was measured within specific subregions of the nucleoid based on probe displacements within macrodomain masks and within nucleoid core and periphery segments.

Macrodomain masks were generated from heatmaps of tracked fluorescent loci using the *kde* function in MATLAB (2024a) to obtain smooth boundaries. Each mask was defined by selecting the grid bins with the highest localization density such that the total enclosed occurrence accounted for 10% of the total occurrence.

The nucleoid outline was defined by a contour line at $p$% of the maximum average displacement as described in the 'Cell boundary and nucleoid outline determination' section. To delineate the core, a narrower contour line was generated at $(p + a)$%, where $a = 5 – 10$% is an adjustable margin. The region enclosed by the $(p + a)$% contour was defined as the nucleoid core (Fig. 5A bottom, dark blue), and the area between the $p$% and $(p + a)$% contours was defined as the nucleoid periphery (Fig. 5A bottom, light blue).

Displacements of probes falling within each region-specific mask (macrodomains, nucleoid core, or nucleoid periphery) were analyzed using the same fitting and calculation procedure described for the total nucleoid viscosity.

*Bootstrapping and statistics*

To estimate population-level statistics and compare conditions, bootstrapping was performed. In each bootstrap iteration, 80% – 90% of the total cells were randomly sampled with replacement from the pool of all cells that passed the filtering criteria, and all relevant calculations were performed on each resampled set.

This resampling procedure was performed 50 times (100 iterations were used when the result did not converge after 50 iterations) based on testing (Fig. S8). Unexpected artifacts—for example, unreasonable nucleoid boundaries and bad fits—that would otherwise require manual intervention in the automated pipeline, were removed as outliers using the MATLAB (2024a) *isoutlier* function with the *median* method. The mean and 95% confidence interval (CI) of each quantity were computed from the resulting bootstrap distributions.

Suitable statistical tests (Fig. S3) were used to compare different conditions. Two-sample t-tests were applied for comparisons between two conditions, with Welch's correction as indicated. Welch's one-way ANOVA with Games-Howell post hoc testing, which accounts for unequal variance, was used to compare accessibility or viscosity across multiple conditions. A one-way repeated-measures ANOVA followed by a Tukey post hoc test was used to compare the viscosities of macrodomains, since these measurements were obtained from the same nucleoid. For the same reason, a one-sided, null-centered bootstrap hypothesis test was used to compare the effective viscosities of the nucleoid periphery and core in Fig. 5A. The difference was defined as $\Delta\eta = \eta_{\text{periphery,10\%}} - \eta_{\text{core,10\%}}$. The bootstrap distribution was centered on the null by subtracting the mean difference, and the *p*-value was calculated using the finite-sampling correction $p = (k + 1)/(B + 1)$ , where $k$ is the number of null-centered differences greater than the observed mean difference and $B$ is the number of bootstrap iterations.

Parameters mentioned in the analysis pipeline can be found in README files in our Github code repository and our Zenodo repository for sample data.

*Adding motion blur and localization error for sensitivity testing*

To assess the sensitivity of effective nucleoid viscosity to motion blur and localization error, we incorporated both effects into trajectories from the 3D Brownian dynamics simulations. Positions were sampled at intervals matching the 40-ms experimental frame time, and motion blur was modeled by taking the averaged original trajectory positions within the temporal window as the new position. After projection onto the *xy* plane, independent Gaussian noise with a standard deviation derived from the experimental localization-fit confidence intervals was added to each coordinate to represent localization error. Displacements between consecutive sampled positions were then calculated, and spatial displacement maps were generated using the same analysis procedure as for simulations without these effects. We found that incorporating reasonable motion blur and localization error has minimal effect on simulated spatial displacement maps and nucleoid step-size distributions (Fig. S9).

*Ensemble mean-squared displacement (MSD) analysis and diffusion coefficient calculation*

Tracks containing more than five observed steps were included. For each trajectory, the time-averaged mean-squared displacement (MSD) was calculated as $MSD(\tau) = \langle [x(t+\tau) - x(t)]^2 + [y(t+\tau) - y(t)^2] \rangle_t$, using observed position pairs separated by the time lag $\tau$ (in this case 40 ms). The ensemble MSD was obtained by averaging individual trajectory MSDs at each lag, with equal weight assigned to each contributing trajectory. Apparent diffusion coefficients were estimated by least-squares fitting of the early-lag of each MSD to the equation $MSD(\tau) = 4D\tau + b$, where $D$ is the diffusion coefficient and $b$ is the offset. Only fits with $R^2 > 0.7$ and $D > 0$ were included in the diffusion coefficient distribution.

*Nanocage fluorescence intensity analysis*

Emitter intensities after background subtraction were quantified using the SMALL-LABS algorithm[2]. To limit photobleaching, we analyzed only the first 10 s of each movie. The brightest nonterminal frame from each trajectory was selected, yielding a histogram of one intensity value per trajectory (Fig. S10). A single gamma distribution and a two-component Gaussian mixture, with the second mean fixed at twice the first, were fit separately to the histogram and compared using the Akaike information criterion (AIC) (Fig. S10).

**Nucleoid Staining and 2D Occupancy Quantification**

Cells were grown as described in the main text and sampled at the 5-h exponential-phase time point. A 1-mL aliquot of culture was incubated with 10 µL of 50 µM SYTOX Orange for 10 min. Cells were then collected by centrifugation and washed twice before imaging. For osmotic shock experiments, treatment was performed as described in the main text for the nanocage tracking experiment. 561-nm laser illumination with a power density of 0.15 kW/cm$^2$ was used for imaging. The nucleoid occupancy was calculated as the ratio of nucleoid area to cell area using the image analysis procedure described in our previous work[3].

**Hi-C Analysis**

Cells used in Hi-C were grown according to the same procedure as described for the single-particle tracking of sfGFP-labeled nanocages. For exponential cultures, cells were first grown in LB supplemented with 50 µg/mL spectinomycin at 37 °C overnight. Next, cells were back-diluted 1:100 in HDA medium containing 0.2% glycerol (m/v) and grown at 37 °C to OD = 0.3. Then, arabinose was added to a concentration of 0.08% for nanocage induction. After 5 h, the culture was again back-diluted 1:100 into HDA medium supplemented with 0.2% glycerol (m/v) and grown at 30 °C. After 5 h of growth, the OD reached 0.25 – 0.3. Cells were harvested for Hi-C. For antibiotic treatment, 25 μg/mL rifampicin (CAS 13292-46-1) or 50 μg/mL chloramphenicol (CAS 56-75-7) was added to cultures for 30 minutes before cell collection.

For 24 h cultures, cells were first grown in LB supplemented with 50 μg/mL spectinomycin at 37 °C overnight. Next, cells were back-diluted 1:100 to HDA medium containing 0.2% glycerol (m/v) and grown at 30 °C until the OD reached 0.15 – 0.20. Then, arabinose was added to a concentration of 0.0013% to express the nanocage. This time is designated time 0. After 24 h, cells were harvested for Hi-C. For antibiotic treatment, 25 μg/mL rifampicin (CAS 13292-46-1) or 50 μg/mL chloramphenicol (CAS 56-75-7) was added to cultures for 30 minutes before cell collection.

The Hi-C procedure was carried out as described previously[3]. Specifically, the collected cells were crosslinked with 10% formaldehyde (Sigma F8775) at room temperature for 45 min, then quenched with 125 mM glycine. Cells were lysed using Ready-Lyse Lysozyme (Epicentre, R1802M) and treated with 0.5% SDS. Solubilised chromatin was digested with Sau3AI for 2 h at 37 °C. The digested ends were filled in with Klenow and Biotin-14-dATP, dGTP, dCTP, dTTP. The products were ligated with T4 DNA ligase at 16 °C for about 20 h. Crosslinks were reversed at 65°C for about 20 h in the presence of EDTA, proteinase K, and 0.5% SDS. The DNA was then extracted twice with 25:24:1 phenol/chloroform/isoamyl alcohol (PCI), precipitated with ethanol, and resuspended in 20 μL of 0.1× TE buffer (pH = 8.0). Biotin from non-ligated ends was removed using T4 polymerase (4 h at 20 °C), followed by extraction with PCI. The DNA was then sheared by sonication for 12 min with 20% amplitude using a Qsonica Q800R2 sonicator. The sheared DNA was used for library preparation with the NEBNext UltraII kit (E7645). Biotinylated DNA fragments were purified using 10 μL streptavidin beads. DNA-bound beads were used for polymerase chain reaction (PCR) in a 50 μL reaction for 14 cycles. PCR products were purified using Ampure beads (Beckman, A63881) and sequenced at the Indiana University Center for Genomics and Bioinformatics using Illumina NextSeq500 or NextSeq2000. Paired-end sequencing reads were mapped to the genome of *E. coli* W3110 (NCBI reference sequence GCA_000005845.2 with a deletion from 1398953 to 1438196) using the same pipeline described previously[5]. The *E. coli* W3110 genome was first divided into 461 10-kb bins. Subsequent analysis and visualisation were performed using R scripts. The *x* and *y* axes were rearranged to start at the replication origin, with the terminus region near the centre of the map. The Hi-C contact probability ($P_c$) curves were calculated by averaging the contact frequency for each pair of loci separated by the set distance specified on the *x*-axis.

**Strain and Plasmid Construction**

LAM022 [W3110, pBad42 I3-01-sfgfp, spec] was generated by transforming chemically competent W3110[6] with pLAM003 by heat shock. Transformants were selected for on

LB spectinomycin 100 μg/mL plates. Colony PCR using primers LM006 and LM027 was performed to confirm plasmid presence.

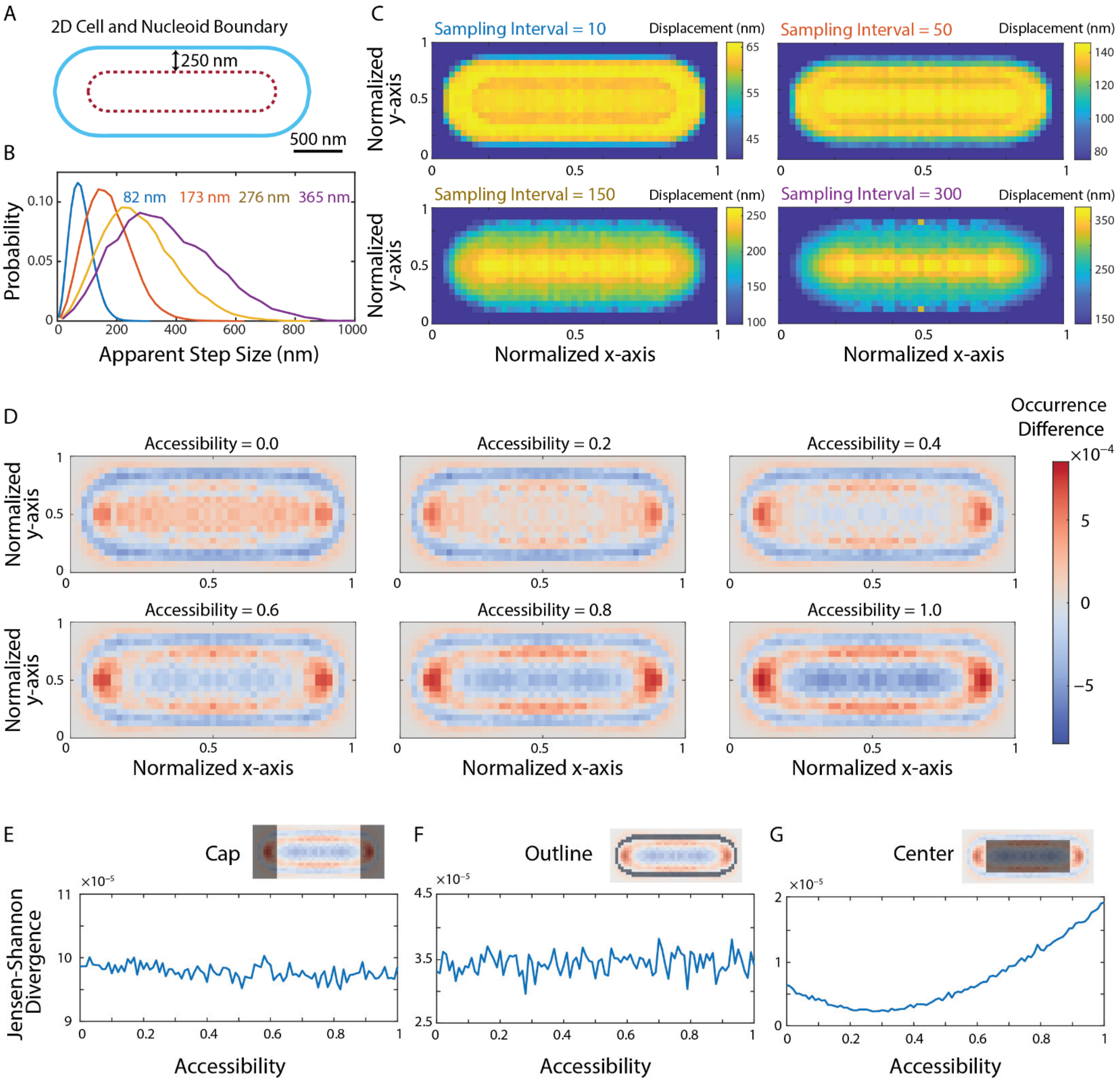


**Figure S1. The effect of computational step size on the displacement map pattern, and comparison of the experimental and simulated localization heatmaps.**

**A**, Example cell and nucleoid geometries used for 3D Brownian dynamics (BD) simulations, shown in two dimensions. Blue line: cell boundary. Red dashed line: nucleoid boundary. Arrow: gap between the nucleoid and the cell inner membrane (250 nm). Simulations of a random walk with step sizes drawn from a Rayleigh distribution about an average step size of 25 nm are sampled at a sampling interval of 10 – 300 steps to reproduce the experimental imaging patterns.

**B**, The apparent step size from the 3D BD simulation increases with sampling interval. Blue: sampling interval = 10, median apparent step size = 82 nm. Orange: sampling interval = 50, median apparent step size = 173 nm. Yellow: sampling interval = 150, median apparent step size = 276 nm. Purple: sampling interval = 300, median apparent step size = 365 nm.

**C**, Displacement maps from 3D BD simulation at sampling intervals of 10, 50, 150, and 300, respectively. When the median apparent step size (**B**) approaches the nucleoid-inner membrane gap size (250 nm in this example), the simulated displacement maps qualitatively reproduce the characteristic pattern observed experimentally (Fig. 1G). This correspondence indicates that geometric confinement imposed by the nucleoid-membrane spacing is a primary driver of the displacement map pattern.

**D**, Pixelwise difference error maps between the experimental localization heatmap (Fig. 1E) and simulated heatmaps generated with different accessibilities.

**E**, Jensen-Shannon (J-S) divergence between the experimental and simulated heatmap within the cap region (dark mask, top right) as a function of accessibility.

**F**, Same as **E**, but within the dark mask outline region.

**G**, Same as **E**, but within the dark mask central region.

We quantified the subregion discrepancies using the J-S divergence rather than the SSIM (Fig. 1) because SSIM is designed for image-level comparisons that rely on global contrast and structural context, while the J-S divergence directly compares probability distributions. Considering that each heatmap is a normalized 2D histogram, the J-S divergence is well-suited for comparing simulated and experimental localization patterns locally. Panels **E** – **G** show that the errors in localization occurrences between experimental and simulated heatmaps within the cap and outline regions are independent of accessibility. As a result, the unmodeled cell shape heterogeneity and cell-pole components do not influence the similarity optimization for accessibility quantification.

Within the projected nucleoid region, the experimental trajectories include both probes diffusing inside the nucleoid and cytoplasmic probes diffusing in the gaps between the nucleoid and the inner membrane. The BD simulations in Fig. S1 A-C show that confinement within these gaps produces large apparent displacements in the projected central region when the net displacement over the 40-ms experimental sampling interval approaches the gap width. Because the probe density is higher in the cytoplasmic gaps than within the nucleoid, the average displacement map is dominated by this cytoplasmic contribution. Thus, the average step size can be greatest in the cell-center region even though probes inside the nucleoid diffuse more slowly. The shorter intra-nucleoid displacements are present in the experimental step-size distribution and are resolved through the two-component fit shown in Fig. 1I.

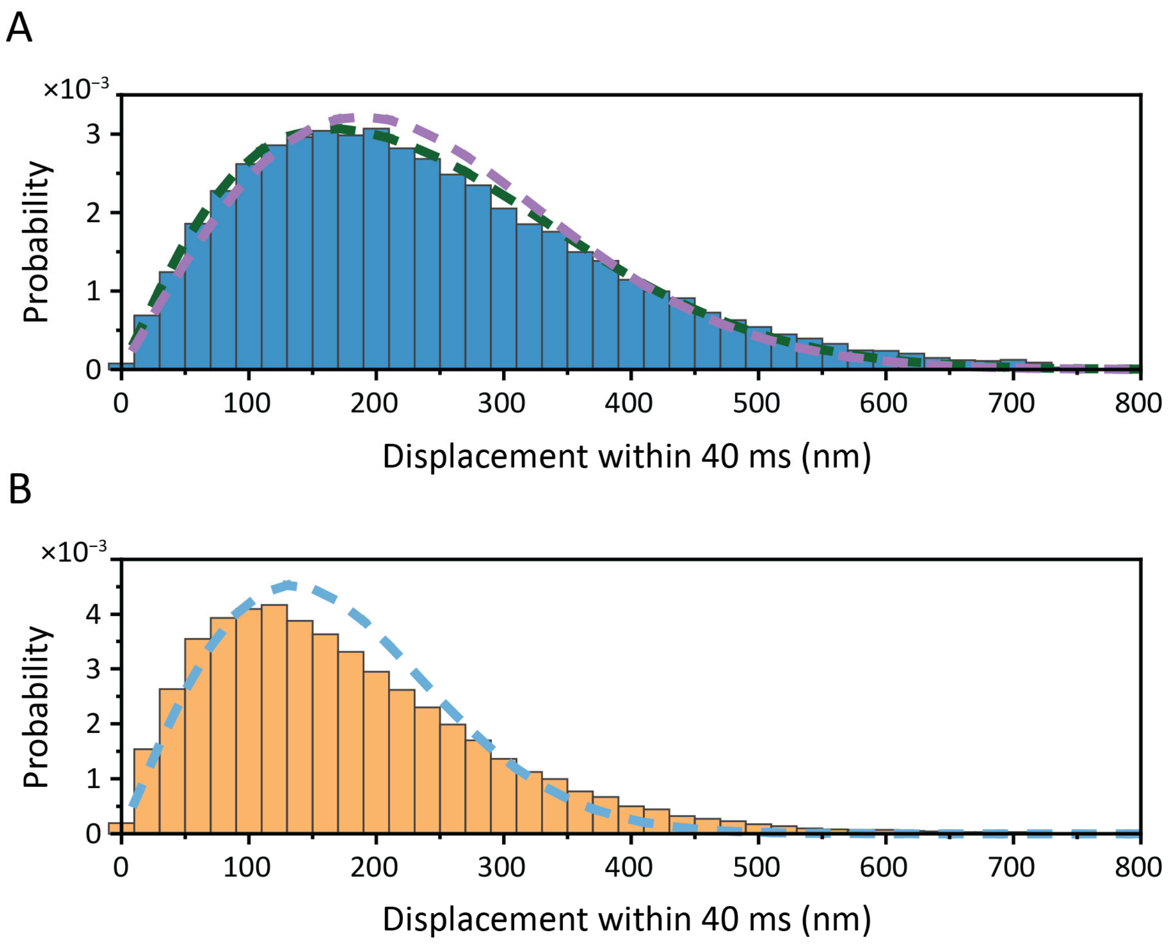


**Figure S2. Rayleigh fits to experimental displacement histograms from the nucleoid and cytoplasm.**

**A,** Histogram of experimental displacements in the nucleoid (middle panel of Fig. 1I) fit to a single Rayleigh distribution (purple dashed line) and a two-component Rayleigh model (green dashed line). A comparison of the single- and two-component Rayleigh fits is shown in Table S6.

**B,** Histogram of experimental displacements in the cytoplasm fit to a single Rayleigh distribution (blue dashed line).

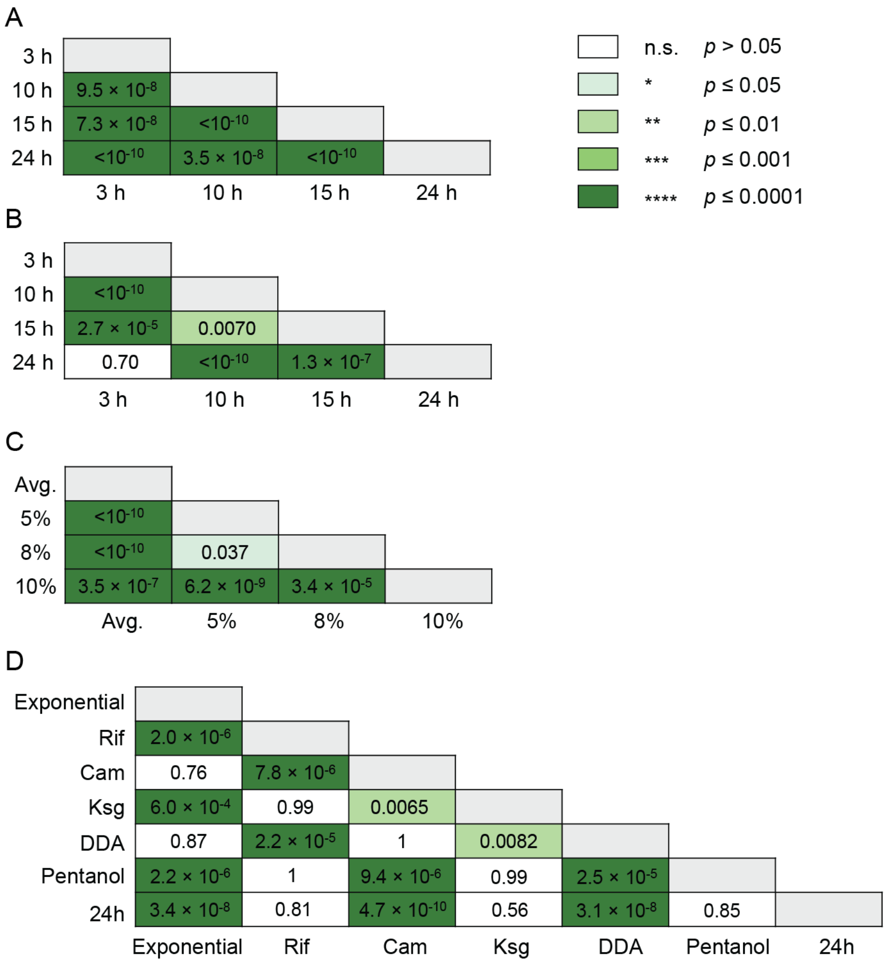


**Figure S3. Statistical tests of pairwise comparisons among different conditions.**

**A**, Apparent nucleoid accessibility comparisons for conditions in Fig. 2B, using a Welch one-way ANOVA followed by a Games-Howell post hoc test.

**B**, Effective nucleoid viscosity comparisons for conditions in Fig. 2B, using a Welch one-way ANOVA followed by a Games-Howell post hoc test.

**C**, Viscosity comparisons between the whole-nucleoid average and the peripheries defined by different margins in Fig. 5A, using a Welch one-way ANOVA followed by a Games-Howell post hoc test.

**D**, Comparisons of periphery-core viscosity differences in Fig. 5B, using a Welch one-way ANOVA followed by a Games-Howell post hoc test.

For all panels, the matrix entries report the *p*-values for each condition pair, and the entry color encodes the significance level as defined in the legend.

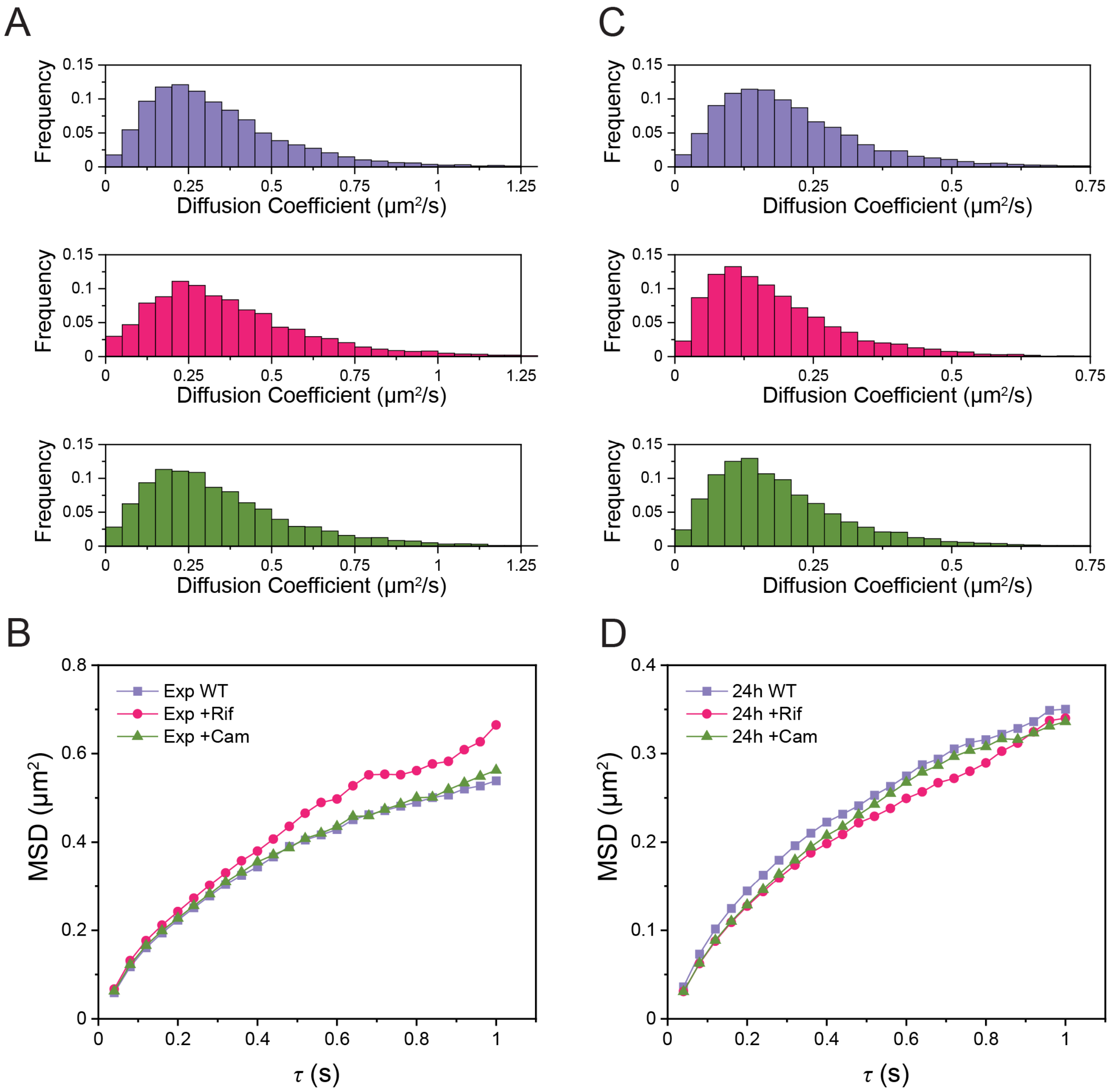


**Figure S4. Effects of transcription and translation inhibition on nanocage diffusion during exponential and stationary phases.**

**A,** Diffusion coefficient distributions for all tracked nanocages during exponential growth in untreated wild-type cells (WT; top, purple), rifampicin-treated cells (Rif; middle, magenta), and chloramphenicol-treated cells (Cam; bottom, green).

**B,** Mean squared displacement (MSD) as a function of lag time ($\tau$) for the conditions in A.

**C,** Diffusion coefficient distributions as in A, but for cells in stationary phase (24 h).

**D,** MSD vs. $\tau$ as in B, but for cells in stationary phase (24 h).

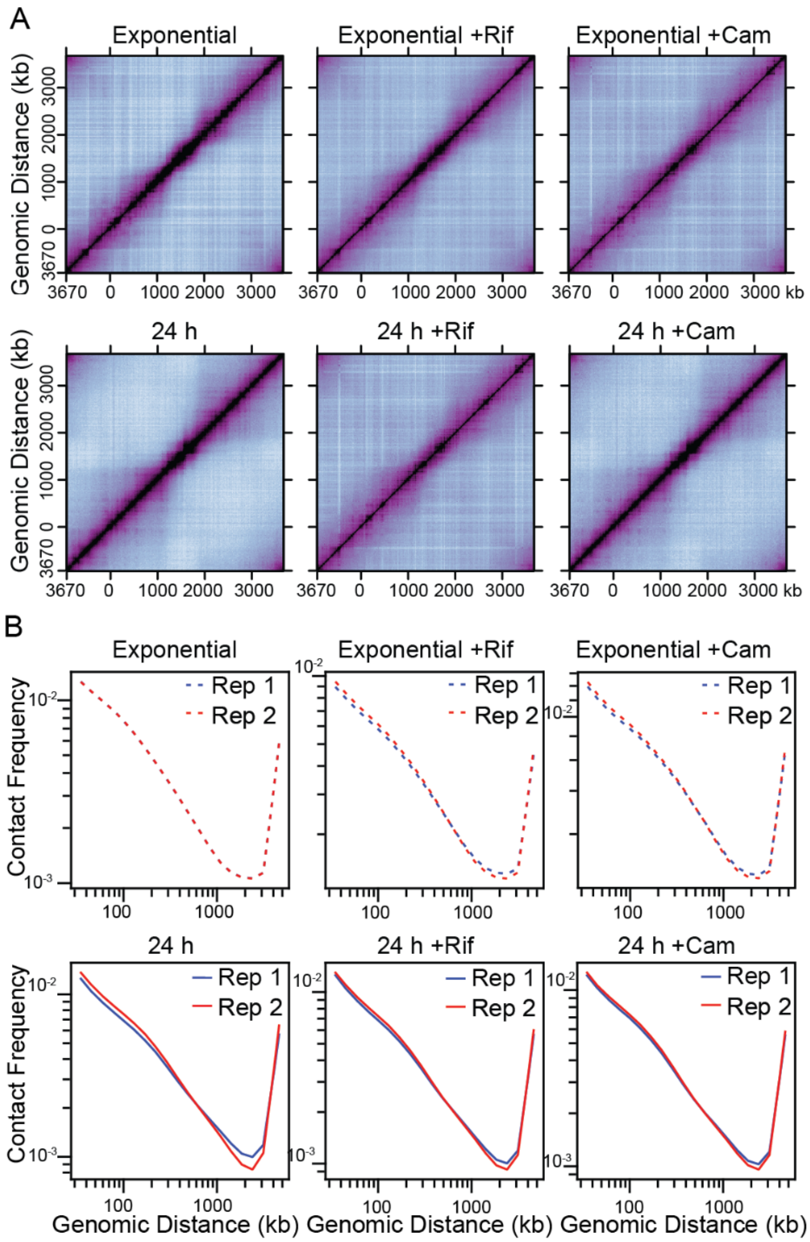


**Figure S5. Comparison of biological replicates of Hi-C experiments.**

**A**, Normalized Hi-C contact maps for *E. coli* without drug treatment, after Rif treatment, or after Cam treatment during exponential phase (row 1) and during the stationary phase (24 h; row 2). These are the second biological replicates of the Hi-C results shown in Fig. 3C – H, using the same color bar shown in Fig. 3K.

**B**, Hi-C contact probability ($P_c$) curves. The *x*-axis indicates the genomic distance, while the *y*-axis shows the averaged contact frequency. Data are presented at a 10-kb resolution. Results from two biological replicates are compared for each of: *E. coli* during the exponential phase without drug, after rifampicin treatment, or after chloramphenicol treatment (row 1), and *E. coli* during the stationary phase (24 h) without drug, after rifampicin treatment, or after chloramphenicol treatment (row 2).

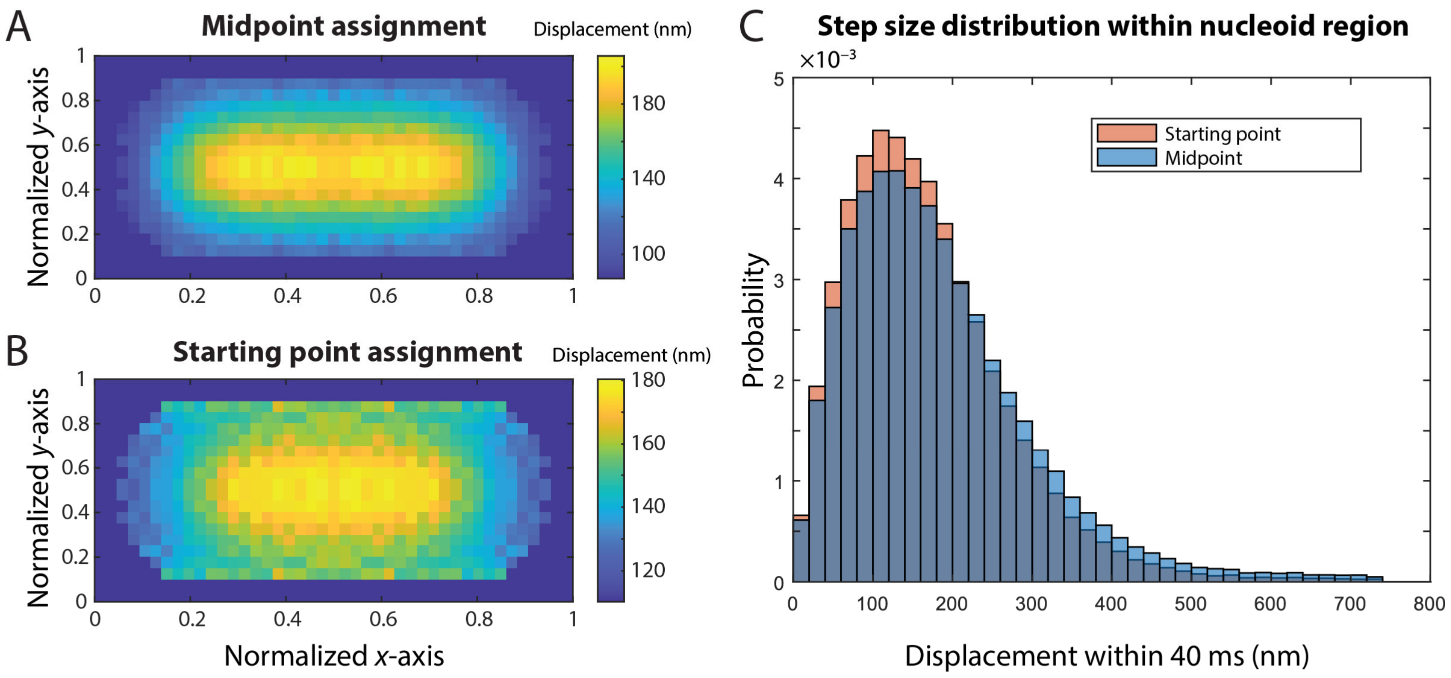


**Figure S6. Sensitivity of simulated displacement maps and nucleoid step-size distributions to step position assignment.**

**A,** Spatial displacement maps from 3D Brownian dynamics simulations constructed by assigning each step to its midpoint.

**B,** Spatial displacement maps from 3D Brownian dynamics simulations constructed by assigning each step to its starting point.

**C,** Nucleoid step-size distributions corresponding to the maps in A and B (blue and orange, respectively).

The larger 24-h WT dataset was used for this sensitivity test, as the higher sample size reduced the pixel-level sampling noise in the comparison.

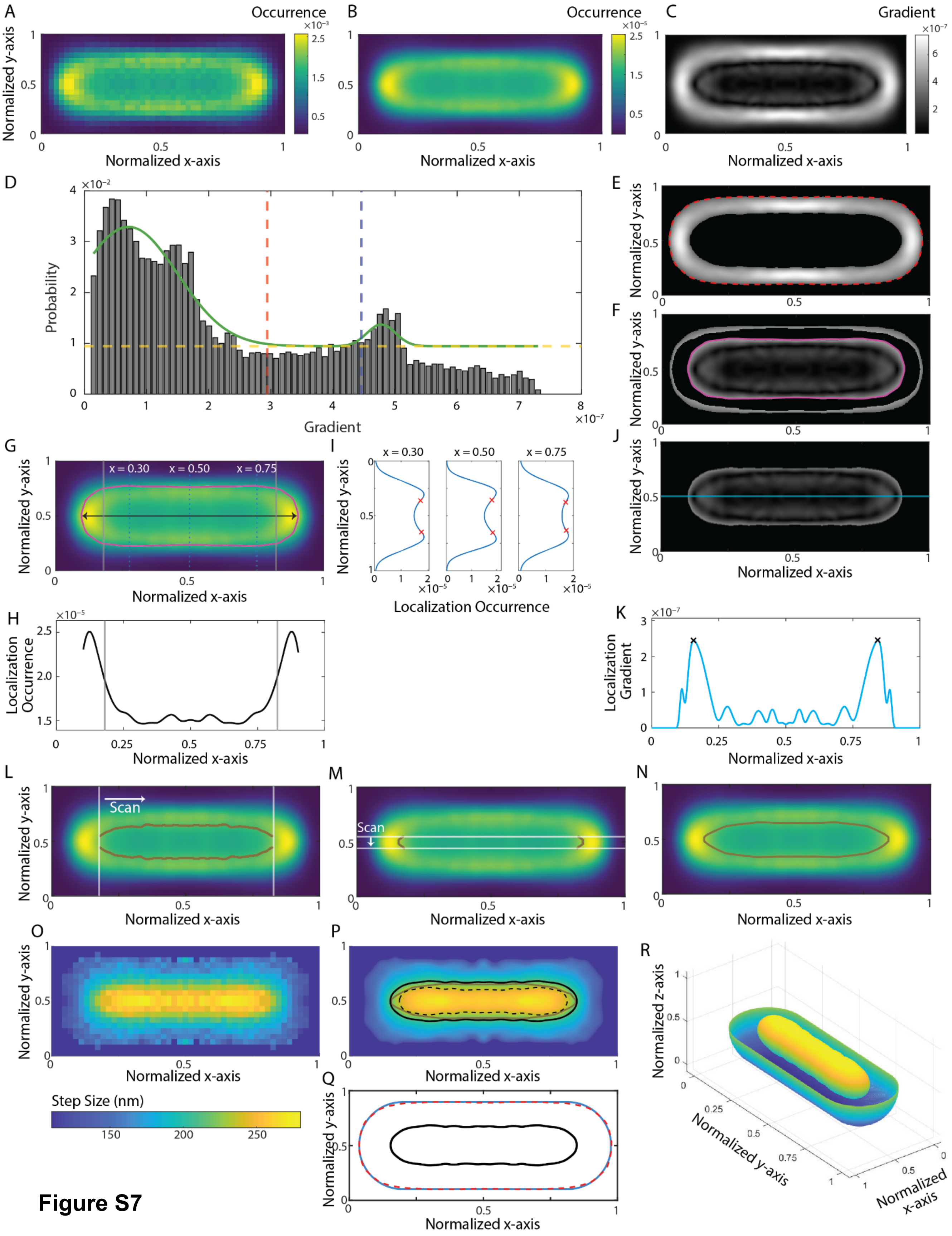
A
B
C
Occurrence
Gradient
Normalized y-axis
Normalized x-axis
D
E
F
Probability
Gradient
G
I
J
x = 0.30
x = 0.50
x = 0.75
Localization Occurrence
H
K
Localization Occurrence
Localization Gradient
L
M
N
Scan
O
P
Q
R
Step Size (nm)
Normalized z-axis

**Figure S7**

**Figure S7. Cell geometry reconstruction.**

**A**, Example experimental nanocage localization heatmap for untreated, exponential-phase cells.

**B**, Smoothed heatmap after 10× interpolation.

**C**, Gradient magnitude map computed from the interpolated heatmap in **B**.

**D**, Histogram of pixelwise gradient values from **C**. Yellow dashed line: baseline level. Green curve: a fit of the values above the baseline to a Gaussian mixture model. The red dashed line indicates the first threshold: the mean plus two standard deviations of the lower Gaussian peak. The blue dashed line indicates the second threshold: the mean minus one standard deviation of the higher Gaussian peak.

**E**, Cell boundary determined from **C** using the first threshold. Pixels in the gradient map above the first threshold are plotted. Colorscale: as in **C**. Red dashed line: region boundary.

**F**, Pixels in the gradient map inside the cell boundary in **E** and with gradient values below the second threshold are plotted. Colorscale: as in **C**. Pink line: Unrefined nucleoid region.

**G**, Interpolated heatmap (**B**) overlaid with the initial nucleoid boundary. Colorscale: as in **B**. Pink line: unrefined nucleoid boundary (**F**). Black arrow: horizontal line at $y = 0.5$ along the long axis within the unrefined nucleoid region used for further analysis in **H**. Gray lines: positions of the steepest change in localization occurrence along the line profile in **H**. Blue dotted lines: $x$ positions used for the example vertical heatmap line profiles **I**.

**H**, Horizontal line profile of localization occurrences along $y = 0.5$. Gray lines: positions of the steepest change.

**I**, Example vertical heatmap line profiles at the indicated $x$ positions. Red crosses: positions of the steepest change between occurrence maxima.

**J**, Pixels in the gradient map inside the unrefined nucleoid region in **F** are plotted. Colorscale: as in **C**. Cyan line: horizontal line at $y = 0.5$ along the long axis used for further analysis in **K**.

**K**, Smoothed gradient profile along cyan line in **J**. Black crosses: gradient maxima.

**L**, Interpolated heatmap (**B**) overlaid with the $x$-scan components of the exclusion region (brown lines) obtained from change points like the red crosses in **I** between the positions of the steepest change from **H** (gray lines).

**M**, Interpolated heatmap (**B**) overlaid with the $y$-scan components of the exclusion region (brown lines) obtained from maxima like the black crosses in **K** in the gap between the brown curves in **L** (white lines) region.

**N**, Interpolated heatmap (**B**) overlaid with the final, refined exclusion region. Brown line: combined exclusion region boundary from **L** and **M**.

**O**, Experimental nanocage displacement map from the same dataset as used for the heatmap in **A**.

**P**, The interpolated displacement map overlaid with the final, refined exclusion region boundary identified in **N** (brown curve). Black solid line: the smallest displacement map contour line that encloses the exclusion region. Black dashed line: the largest displacement map contour line that is fully contained within the exclusion region.

**Q**, The 2D cell boundary identified in **E** (red dashed line) is fit to a rod shape (blue solid line). Black line: the 2D nucleoid boundary defined in **P**.

**R**, 3D reconstructed cell and nucleoid geometries constructed by rotating the 2D outlines in **Q** about the long axis. The upper half of the cell surface is omitted for nucleoid visualization.

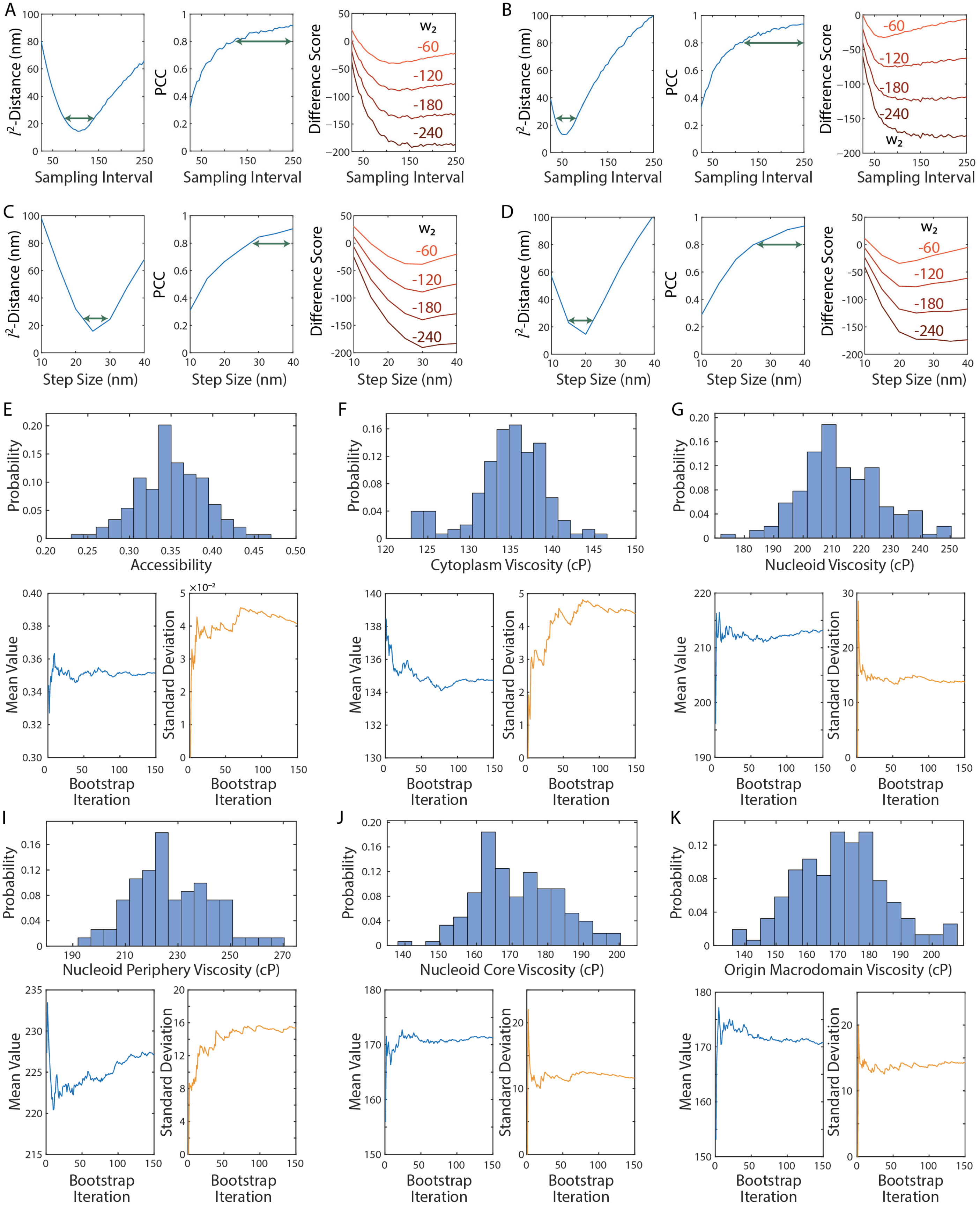
A
B
C
D
E
F
G
I
J
K
$l^2$-Distance (nm)
PCC
Difference Score
Sampling Interval
Step Size (nm)
$w_2$
-60
-120
-180
-240
Probability
Accessibility
Cytoplasm Viscosity (cP)
Nucleoid Viscosity (cP)
Nucleoid Periphery Viscosity (cP)
Nucleoid Core Viscosity (cP)
Origin Macrodomain Viscosity (cP)
Mean Value
Standard Deviation
Bootstrap Iteration

**Figure S8**

**Figure S8. Weight parameter justification and bootstrapping convergence.**

**A**, $l^2$-distance ($d_{L2}$), Pearson correlation coefficient (PCC), and difference score, $S = 0.5 \times d_{L2} + w_2 PCC$, as a function of sampling interval from 3D Brownian dynamics simulations (average step size = 25 nm) within cell and nucleoid geometries reconstructed from experiments on untreated cells at exponential phase. Green arrows: target sampling interval range based on the dip in $l^2$-distances and the largest PCC values (left and middle panels, respectively). $w_2$ values are labeled in the right panel.

**B**, Same as **A**, but using geometries reconstructed from untreated cells at 24 h.

**C**, $l^2$-distance ($d_{L2}$), Pearson correlation coefficient (PCC), and difference score, $S = 0.5 \times d_{L2} + w_2 PCC$, as a function of sampling interval from 3D Brownian dynamics simulations (sampling Interval = 100) within cell and nucleoid geometries reconstructed from experiments on untreated cells at exponential phase. Green arrows: target sampling interval range based on the dip in $l^2$-distances and the largest PCC values (left and middle panels, respectively). $w_2$ values are labeled in the right panel.

**D**, Same as **C**, but using geometries reconstructed from untreated cells at 24 h.

To estimate population-level statistics and compare conditions, bootstrapping was performed. In each bootstrap iteration, 80 – 90% of the total cells were randomly sampled with replacement from the pool of all cells that passed the filtering criteria, and all relevant calculations were performed on each resampled set.

**E**, Apparent nucleoid accessibility. Top: Histogram of accessibility from 150 bootstrap iterations. Bottom: mean value (left) and standard deviation (right) versus number of bootstrap iterations. Both the mean and standard deviation of the bootstrap distribution are used to assess bootstrap convergence.

**F**, Same as **E**, but for the effective cytoplasm viscosity.

**G**, Same as **E**, but for the effective nucleoid viscosity.

**H**, Same as **E**, but for the effective nucleoid periphery viscosity.

**I**, Same as **E**, but for the effective nucleoid core viscosity.

**J**, Same as **E**, but for the effective Ori macrodomain viscosity.

The Shapiro-Wilk normality tests for all bootstrap distributions are presented in Table S12.

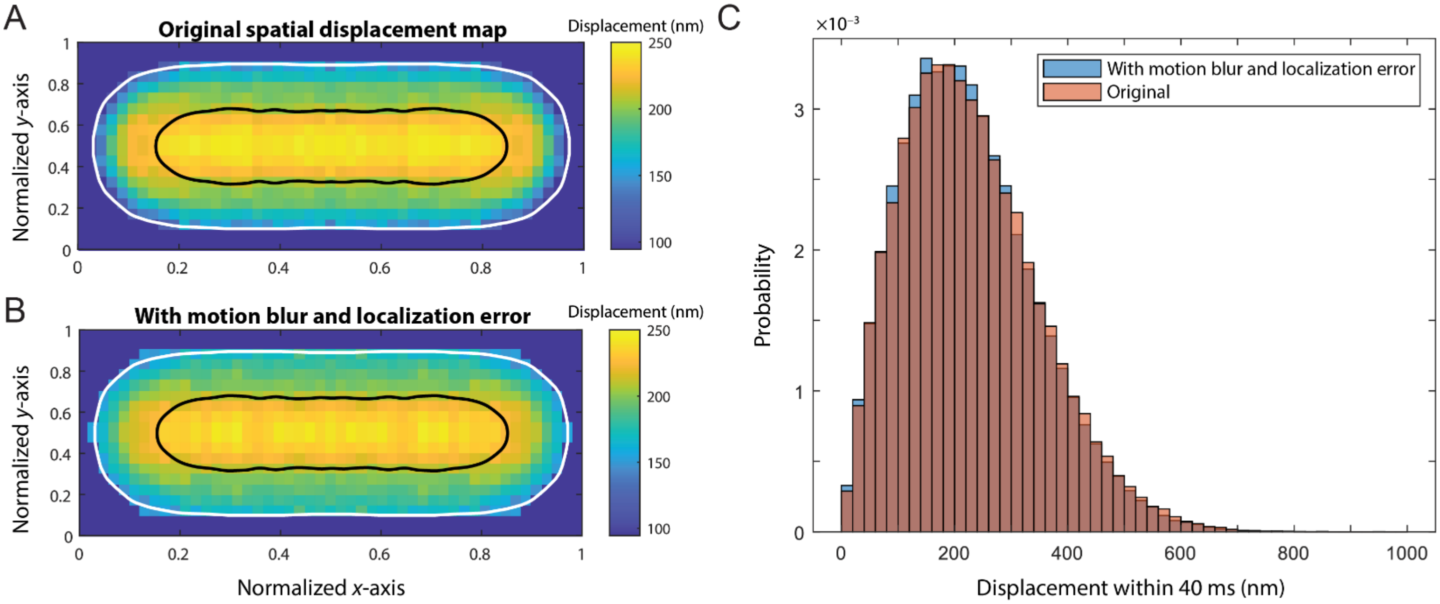


**Figure S9. Localization error and motion blur have minimal effect on simulated spatial displacement maps and nucleoid step-size distributions.**

**A,** Spatial displacement maps from the original 3D Brownian dynamics simulation (Fig. 1G, right). White and black outlines indicate the cell and nucleoid boundaries, respectively.

**B,** Spatial displacement map after incorporating motion blur and localization error matching the tracking experiments. White and black outlines indicate the cell and nucleoid boundaries, respectively.

**C,** Nucleoid step-size distributions from the original simulations (orange) and simulations incorporating motion blur and localization error (blue).

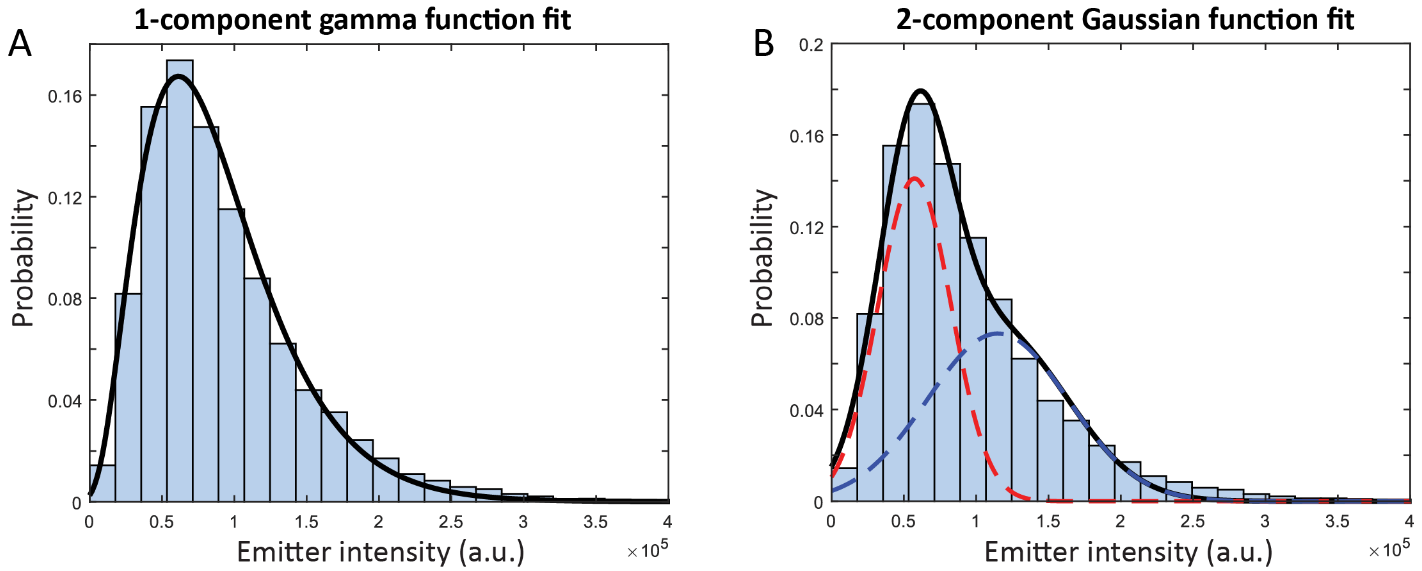


**Figure S10. Fluorescence intensity distribution of tracked nanocages.**

**A,** Single gamma function fit (solid black line) to the fluorescence intensity distribution of tracked nanocages

**B,** Two-component Gaussian mixture fit to the same distribution. The solid black line indicates the overall fit; the dashed red and blue lines indicate the individual Gaussian components.

For each nanocage trajectory, we recorded the brightest value and plotted the distribution of these intensities. The brightness distribution is unimodal and right-skewed, with no resolved secondary peak at values around twice the modal brightness. The tail could reflect differences in focal position, excitation intensity, or the number of mature and unbleached sfGFP molecules per cage. Selecting the brightest frame may also preferentially select for upward fluctuations arising from photon-counting and camera noise. We used a gamma distribution as a descriptive model because it accommodates positive, right-skewed brightness values. A single Gaussian is symmetric and permits negative brightness values, whereas a two-component Gaussian mixture can approximate a single skewed distribution without establishing the existence of two distinct stoichiometric species. The single gamma model also provided a better fit than the two-component Gaussian mixture, with lower AIC (667 versus 3790) and RMSE ($1.2 \times 10^{-3}$ versus $3.7 \times 10^{-3}$).

### SI Tables

**Table S1.** Strains used in this study.

| Strain | Genotype | Reference | Figures |
|---|---|---|---|
| LAM022 | W3110, pBad42 I3-01-sfgfp, spec (Nanocage) | This study | 1ABJ, 2ABCD, 3A-J, 4C, 5AB, S3 |
| cWX2565 | W3110 *yidA(ori) PdnaA ygfp-ParBMT1-parSMT1 FRT-kan-FRT* | McCarthy. et al.[3] | 4ABC |
| cWX2610 | W3110, *ydeA (ter) PdnaA ygfp-ParBMT1-parSMT1 FRT-kan-FRT* | McCarthy. et al.[3] | 4ABC |
| Strains used to build strains | | | |
| W3110 | *E. coli* K-12 strain W3110 ((Migula) Castellani and chalmers) *F-, λ-, IN(rrnD-rrnE)1, rph-1* | ATCC27325 | |

**Table S2.** Plasmids used in this study.

| Plasmid | Description | Reference |
|---|---|---|
| pLAM003 | *pBad42 I3-01-sfgfp, spec* | McCarthy. et al.[3] |

**Table S3.** Oligonucleotides used in this study.

| Oligos | Sequence | Plasmid |
|---|---|---|
| LM006 | GCGTTCTGATTTAATCTGTATCAGG | pLAM003 |
| LM027 | CACACTTTGCTATGCCATAGC | pLAM003 |

**Table S4.** NGS samples used in this study (reviewer token: aduhkwgkhdcbhin).

| Sample Name | Figure | Reference | Identifier |
|---|---|---|---|
| HiC_cWX3118_LAM022_Exp_Azure10percent_Sau_rep1 | 3C, 3I, S3B | This study | GSM9382719 |
| HiC_cWX3118_LAM022_Exp_Azure10percent_Sau_rep2 | S3A, S3B | This study | GSM9382720 |
| HiC_cWX3118_LAM022_rif_Exp_Azure10percent_Sau_rep1 | 3D, 3I, S3B | This study | GSM9382721 |
| HiC_cWX3118_LAM022_rif_Exp_Azure10percent_Sau_rep2 | S3A, S3B | This study | GSM9382722 |
| HiC_cWX3118_LAM022_cm_Exp_Azure10percent_Sau_rep1 | 3E, 3I, S3B | This study | GSM9382723 |
| HiC_cWX3118_LAM022_cm_Exp_Azure10percent_Sau_rep2 | S3A, S3B | This study | GSM9382724 |
| HiC_cWX3118_LAM022_24h_Azure10percent_Sau_rep1 | 3F, 3J, S2B | This study | GSM9382713 |
| HiC_cWX3118_LAM022_24h_Azure10percent_Sau_rep2 | S3A, S3B | This study | GSM9382716 |
| HiC_cWX3118_LAM022_rif_24h_Azure10percent_Sau_rep1 | 3G, 3J, S3B | This study | GSM9382714 |
| HiC_cWX3118_LAM022_rif_24h_Azure10percent_Sau_rep2 | S3A, S3B | This study | GSM9382718 |
| HiC_cWX3118_LAM022_cm_24h_Azure10percent_Sau_rep1 | 3H, 3J, S3B | This study | GSM9382715 |
| HiC_cWX3118_LAM022_cm_24h_Azure10percent_Sau_rep2 | S3A, S3B | This study | GSM9382717 |

**Table S5.** Antibiotic and chemical conditions*.

| Chemical | CAS No. | Concentration | Usage | Manufacturer |
|---|---|---|---|---|
| Sodium chloride | 7647-14-5 | 200 mM | Figure 1J | Sigma-Aldrich |
| Rifampicin | 13292-46-1 | 25 µg/mL | Figure 3A, 5B | Sigma-Aldrich |
| Chloramphenicol | 56-75-7 | 50 µg/mL (imaging), 20 µg/mL (culture) | Figure 3B, 5B, cell culture | Sigma-Aldrich |
| Kasugamycin hydrochloride | 19408-46-9 | 1 mg/mL | Figure 5B | Sigma-Aldrich |
| Dodecylamine hydrochloride | 929-73-7 | 20 µM | Figure 5B | Fisher Scientific |
| 1-pentanol | 71-41-0 | 10 mM | Figure 5B | Sigma-Aldrich |
| Kanamycin | 25389-94-0 | 20 µg/mL | Cell culture | Fisher Bioreagents |
| Spectinomycin | 22189-32-8 | 100 µg/mL | Cell culture | MP Biomedicals |
| SYTOX Orange | N/A | 500 nM | DNA staining | Thermo Fisher Scientific |

*We prepared rifampicin as a 50 mg/mL stock in DMSO and diluted it 1:2000 to a working concentration of 25 µg/mL, resulting in a final DMSO concentration of 0.05% (v/v). Chloramphenicol was prepared as a 25 mg/mL stock in ethanol and diluted 1:500 to a working concentration of 50 µg/mL, resulting in a final ethanol concentration of 0.2% (v/v). All other compounds were dissolved in water.

**Table S6.** Comparison of single- and two-component Rayleigh fits to the experimental displacement histogram. The experimental displacement histogram (Fig. 1I middle panel) fit to a single Rayleigh distribution (one fit parameter) was compared to a fit to a two-component Rayleigh distribution (three fit parameters). We compared the Residual sum of squares (RSS), the Akaike Information Criterion (AIC), and the Bayesian Information Criterion (BIC). Both fits used forty bins of width 20 nm (Fig. S2).

| Comparison | Single Rayleigh | Two-Component Rayleigh |
|---|---|---|
| RSS | $1.04 \times 10^{-6}$ | $2.19 \times 10^{-7}$ |
| AIC | −581.21 | −639.44 |
| BIC | −577.83 | −632.68 |

**Table S7.** Nucleoid effective viscosity calculated for the optimal parameter pair and neighboring parameter pairs. Each cell lists the parameter pair (simulation step size in nm, sampling interval) on the first line, and the corresponding effective viscosity with the 95% confidence interval reported as [lower bound, upper bound] in cP. The optimal parameter pair is highlighted in bold.

| | | |
|---|---|---|
| | 20,135<br>Cannot be fit, large error | |
| 25,130<br>252 cP [198, 346] | **25, 135**<br>**186 cP [162,219]** | 25,140<br>162 cP [146, 181] |
| | 30,135<br>121 cP [115, 127] | |

**Table S8.** Nucleoid area ratios measured by DNA staining in control and NaCl-treated cells. The nucleoid area ratio decreased significantly from (69.8 ± 0.6%) in control cells to (65.0 ± 0.5%) following NaCl treatment (mean ± 95% confidence interval; Welch's two-sample t-test, $p = 1.7 \times 10^{-9}$). This direct DNA staining measurement therefore supports nucleoid compaction under the osmotic treatment condition.

| Condition | Number of Cells | Nucleoid Area Ratio (%) | 95% CI (%) |
|---|---|---|---|
| Control | 526 | 69.8 | 1.2 |
| NaCl | 489 | 65.0 | 1.0 |

**Table S9.** Cytoplasm effective viscosity at different growth time points. Growth time points correspond to those shown in Fig. 2A-B. Values are reported as bootstrap means with 95% confidence intervals estimated from 50 bootstrap resamples.

| Growth Timepoint (h) | Cytoplasm Effective Viscosity (cP) |
|---|---|
| 5 | 82 ± 0.9 |
| 10 | 82 ± 2 |
| 15 | 100 ± 1 |
| 24 | 135 ± 0.8 |

**Table S10.** Nucleoid and cytoplasm effective viscosities and apparent accessibilities under the conditions shown in Fig. 3A-B. Values are reported as bootstrap means with 95% confidence intervals estimated from 50 bootstrap resamples.

| Condition | Nucleoid Effective Viscosity (cP) | Cytoplasm Effective Viscosity (cP) | Apparent Accessibility (%) |
|---|---|---|---|
| Exponential WT | 213 ± 18 | 82 ± 0.6 | 28 ± 1.5 |
| Exponential +Rif | 133 ± 10.5 | 67 ± 1.6 | 44 ± 4 |
| Exponential +Cam | 215 ± 11.4 | 74 ± 0.9 | 20 ± 2 |
| 24 h WT | 223 ± 6 | 135 ± 0.8 | 36 ± 1.2 |
| 24 h +Rif | 281 ± 4.7 | 172 ± 1.8 | 32 ± 1.1 |
| 24 h +Cam | 324 ± 5.5 | 163 ± 1.1 | 19 ± 0.7 |

**Table S11.** Effective viscosities of the nucleoid core, nucleoid periphery, and cytoplasm under the conditions shown in Fig. 5B. Values are reported as bootstrap means with 95% confidence intervals estimated from 50 bootstrap resamples.

| Condition | Core Effective Viscosity (cP) | Periphery Effective Viscosity (cP) | Cytoplasm Effective Viscosity (cP) |
|---|---|---|---|
| Exponential | 192 ± 20 | 176 ± 8 | 82 ± 0.9 |
| Exponential +Rif | 106 ± 5 | 150 ± 10 | 69 ± 1 |
| Exponential +Cam | 186 ± 11 | 200 ± 5 | 74 ± 0.7 |
| Exponential +Ksg | 141 ± 10 | 186 ± 15 | 74 ± 0.9 |
| Exponential +DDA | 358 ± 27 | 386 ± 16 | 124 ± 2 |
| Exponential +Pentanol | 119 ± 6 | 170 ± 8 | 81 ± 0.8 |
| 24 h | 179 ± 3 | 263 ± 6 | 135 ± 0.8 |

**Table S12.** Shapiro-Wilk normality tests for the bootstrap distributions shown in Fig. S8. At a significance threshold of $p < 0.05$, only the cytoplasmic viscosity (Fig. S8F) showed a statistically significant departure from normality, and this distribution was in fact unimodal and approximately bell-shaped by visual inspection; the rejection appears to be driven primarily by an excess of values in the lowest-viscosity bin rather than by strong skewness, multimodality, or a distinct second population.

| Property | *p*-value |
|---|---|
| Accessibility | 0.99 |
| Cytoplasm Viscosity | $8.2 \times 10^{-5}$ |
| Nucleoid Viscosity | 0.26 |
| Nucleoid Periphery Viscosity | 0.14 |
| Nucleoid Core Viscosity | 0.17 |
| Origin Macrodomain Viscosity | 0.82 |
